# Exploring mechanisms that affect coral cooperation: symbiont transmission mode, cell density and community composition

**DOI:** 10.1101/067322

**Authors:** Carly D Kenkel, Line K Bay

**Affiliations:** Department of Biological Sciences, University of Southern California, 3616 Trousdale Parkway, Los Angeles, CA 90089, USA; Australian Institute of Marine Science, PMB No 3, Townsville MC, Queensland 4810, Australia

**Author notes:** DATA ARCHIVAL LOCATION: Raw amplicon sequencing data have been uploaded to NCBI’s SRA: PRJNA338365. All R scripts and input files have been included as supplementary files for review and will be uploaded to Zenodo upon publication DOI: 10.5281/zenodo.1208684.

## Abstract

The coral symbiosis is the linchpin of the reef ecosystem, yet the mechanisms that promote and maintain cooperation between hosts and symbionts have not been fully resolved. We used a phylogenetically controlled design to investigate the role of vertical symbiont transmission, an evolutionary mechanism in which symbionts are inherited directly from parents, predicted to enhance cooperation and holobiont fitness. Six species of coral, three vertical transmitters and their closest horizontally transmitting relatives, which exhibit environmental acquisition of symbionts, were fragmented and subjected to a two-week thermal stress experiment. Symbiont cell density, photosynthetic function and translocation of photosynthetically fixed carbon between symbionts and hosts were quantified to assess changes in physiological performance and cooperation. All species exhibited similar decreases in symbiont cell density and net photosynthesis in response to elevated temperature, consistent with the onset of bleaching. Yet baseline cooperation, i.e. translocation of photosynthate, in ambient conditions and the reduction in cooperation in response to elevated temperature differed among species. Although *Porites lobata* and *Galaxea acrhelia* did exhibit the highest levels of baseline cooperation, we did not observe universally higher levels of cooperation in vertically transmitting species. *Post hoc* sequencing of the *Symbiodinium* ITS-2 locus was used to investigate the potential role of differences in symbiont community composition. Interestingly, reductions in cooperation at the onset of bleaching tended to be associated with increased symbiont community diversity among coral species. The theoretical benefits of evolving vertical transmission are based on the underlying assumption that the host-symbiont relationship becomes genetically uniform, thereby reducing competition among symbionts. Taken together, our results suggest that it may not be vertical transmission *per se* that influences host-symbiont cooperation, but genetic uniformity of the symbiont community, although additional work is needed to test this hypothesis.

## INTRODUCTION

Cooperation between species has played a fundamental role in the evolution and diversification of life (Friesen & Jones 2013; Kiers & West 2015; Maynard Smith & Szathmary 1995; Moran 2006). In the case of reef-building corals, the intracellular symbiosis between dinoflagellates in the genus *Symbiodinium* and a calcifying Cnidarian host forms the basis of one of the most biodiverse and productive ecosystems on the planet (Hatcher 1988; Knowlton et al. 2010). The process of host calcification, which builds the three dimensional structure of the reef, is largely powered by symbiont primary productivity (Roth 2014). However, climate change and other anthropogenic processes threaten reefs because of the sensitivity of the coral-dinoflagellate symbiosis to environmental stress (Hoegh-Guldberg et al. 2007; Hughes et al. 2003), indicating that host-symbiont cooperation is not stable over ecological timescales.

Recent work has suggested that a transition to parasitism may precipitate the breakdown of the host-symbiont relationship (Baker et al. 2018), but this is not a unique feature of the coral symbiosis. Across taxa, mutualisms are better defined as a spectrum that ranges from negative parasitic interactions to mutually beneficial symbioses, both within the context of a focal inter-species interaction and when comparing relationships across taxa (Doebeli & Knowlton 1998; Lesser et al. 2013; Nowak et al. 1994; Sachs et al. 2011). Less well-resolved, particularly for the coral-*Symbiodinium* symbiosis, are the mechanisms that promote and maintain positive interactions between partners (Lesser et al. 2013; Sachs & Wilcox 2006).

One major factor predicted to influence levels of cooperation is the mode of symbiont transmission (Anderson & May 1982; Ebert & Bull 2003). In corals as in other symbioses, two transmission modes predominate: symbionts can be acquired horizontally from the local environment, usually during a defined larval stage, or vertically from parents, typically through the maternal germ line (reviewed in (Bright & Bulgheresi 2010). Virulence theory predicts that horizontal transmission allows symbionts to adopt selfish strategies, potentially harmful to the host (Bull 1994). A transition from horizontal to vertical transmission is predicted to align the reproductive interests of partners (via partner-fidelity feedback *sensu* (Sachs et al. 2004) and optimize resource sharing to maximize holobiont (the combination of host and symbiont) fitness (Ebert 2013; Frank 1994; Herre et al. 1999).

Experimental manipulations of transmission mode in other systems have provided empirical support for a reduction in pathogen virulence under enforced vertical transmission scenarios (Bull et al. 1991; Dusi et al. 2015; Sachs & Wilcox 2006; Stewart et al. 2005). Bacteriophages forced into vertical transmission evolved lower virulence and lost the capacity to transmit horizontally (Bull et al. 1991). Similarly, *Symbiodinium microadriaticum* under an experimentally enforced horizontal transmission regime proliferated faster within their *Cassiopea* jellyfish hosts while reducing host reproduction and growth (Sachs & Wilcox 2006). However, studies attempting to quantify the evolutionary consequences of natural shifts in transmission mode remain rare, with Herre’s demonstration of a negative relationship between vertical transmission and virulence in nematodes that parasitize fig wasps a notable exception (Herre 1993).

Reef-building corals are a potential system in which to study naturally occurring transitions in transmission mode in a mutualistic symbiosis. Corals (Cnidaria: Anthozoa: Scleractinia) are colonial animals that harbour intracellular populations of dinoflagellate algae in the genus *Symbiodinium*. This symbiosis is considered obligate as the breakdown of the relationship between host animals and their intracellular *Symbiodinium* algae, commonly known as coral bleaching, has major fitness consequences for both partners, and can be lethal (reviewed in (Brown 1997). This inter-species partnership is ancient (evolved ∼ 250 MYA (Stanley & Swart 1995), prolific (600+ coral species worldwide (Daly et al. 2007), and constitutes the foundation of one of the most bio-diverse ecosystems on the planet. The majority of coral species (∼85%) acquire their *Symbiodinium* horizontally from the local environment in each generation (Harrison & Wallace 1990). However, vertical transmission has independently evolved at least four times, such that both transmission strategies can be exhibited by different coral species within the same genus (Baird et al. 2009b; Hartmann et al. 2017).

We compared physiological components of cooperation and fitness proxies between horizontal and vertical transmitters in a phylogenetically controlled design using three pairs of related coral species exhibiting different strategies: (1) *Galaxea acrhelia* (vertical transmitter, VT) and *G. astreata* (horizontal transmitter, HT); (2) *Porites lobata* (VT) *and Goniopora columna* (HT); (3) *Montipora aequituberculata* (VT) *and Acropora millepora* (HT). Species comparisons were drawn from the same or sister genera and replicate comparisons from more distantly related clades (Hartmann et al. 2017). Species also represented a diversity of reproductive modes (e.g. broadcast spawner, brooder), sexual systems (e.g. hermaphroditic, gonochoric), and morphologies (e.g. massive, branching), and host different subclades of *Symbiodinium* (Tonk et al. 2013), such that mode of symbiont transmission was the only consistent difference between pairs (Baird et al. 2009b; Franklin et al. 2012; Kerr et al. 2011). We quantified changes in symbiont cell density, photosynthetic function and translocation of photosynthetically fixed carbon between symbionts and hosts. We defined cooperation as the proportion of photosynthetically fixed carbon translocated to the host, while the degree of symbiont parasitism was calculated as the difference in the proportion of photosynthetically fixed carbon translocated to hosts between control and heat-treated samples at the end of the experiment, *sensu* Baker et al. (2018). Bleaching, or the reduction in symbiont density in response to sustained thermal stress was used as a proxy for holobiont fitness. While we observed differences in host-symbiont cooperation, both at a baseline level and during the onset of bleaching, vertically transmitting species did not exhibit universally elevated levels of cooperation. Additional *post hoc* analysis of *Symbiodinium* ITS-2 diversity among coral species was therefore used to investigate whether symbiont community composition could better explain physiological trait patterns. Symbiont community composition did not explain a significant portion of the variation in physiological components of cooperation or fitness proxies; however, diversity tended to be associated with the degree of symbiont parasitism at the onset of bleaching, suggesting that reduced genetic diversity of symbionts, rather than vertical transmission *per se*, may influence host-symbiont cooperation.

## MATERIALS and METHODS

### Coral Collection and Acclimation

Fragments from sixty unique coral colonies, ∼ 20 cm in diameter, were collected from reefs on the Central GBR from the 8-22 April 2015 under the Great Barrier Reef Marine Park Authority permits G12/35236.1 and G14/37318.1, prioritizing collection of focal species pairs by transmission mode from the same reef environment, including location and depth. Ten corals of each species, *Galaxea acrhelia* and *G. astreata* were collected from Davies Reef (18°49.816’, 147°37.888’, 11 April 2015), ten corals each of *Acropora millepora* and *Montipora aequituberculata* were collected from Pelorus Island (18°33.358’, 146°30.276’, 18 April 2015) and ten corals each of *Goniopora columna* and *Porites lobata* were collected from Pandora Reef (18°48.778’, 146°25.593’, 21 April 2015) from depths of <10m across all reefs, and from the same depth within each site. Corals were transported to the Australian Institute of Marine Science Sea Simulator facility and placed in shaded outdoor holding tanks with 0.2 µM filtered flow-through seawater (FSW, 27°C, 150 µmol quanta m^−2^ s^−1^). Each individual coral colony was further cut into 6 replicates using a diamond blade band saw and these fragments were mounted on aragonite plugs using either super-glue or marine epoxy. On 1 May 2015, fragments were moved into indoor experimental aquaria, which consisted of 12 50-L treatment tanks fitted with 3.5-watt Turbelle nanostream 6015 pumps (Tunze, Germany) with flow-through filtered seawater (FSW, ∼25 liters/hour) at 27°C with Hydra 52 lights (Aqua Illumination, C2 Development, Inc., USA) on a 12:12 light/dark cycle mimicking natural irradiance patterns, peaking at 130-160 µmol quanta m^−2^ s^−1^ at “midday”, which actually occurred at 10:30 GMT+10. Tanks, coral racks and plugs were cleaned daily. Beginning on 7 May, at 20 mins before dark (16:30 GMT +10) every day, corals were fed *Artemia* naupli at a density of 1-1.5 naupli/5ml and rotifers at a density of 1-3/ml.

### Experimental Conditions and Physiological Trait Measurements

Beginning on 28 May 2015, effective quantum yield of *Symbiodinium* photosystem II (EQY) was measured daily for all experimental fragments using a pulse amplitude modulated fluorometer (diving-PAM, Walz) fitted with a plastic fibre optic cable (Fig. S1).

Measurements were made using factory settings with a measuring intensity of 12 and a gain of 5 and taken at peak light intensity (between 09:30 – 1130 GMT +10). EQY values were used to guide the timing of the final sampling time point (Fig. S2), where a decline reflects an impact on the photosynthetic condition of the *Symbiodinium* (Ralph et al. 2005) because we aimed to target the onset of the coral bleaching response rather than the end-point.

On 31 May 2015, temperatures in the heat treatment tanks were increased at a rate of 1°C per day until temperatures reached 31°C (day 4, Fig S3). Sample time points occurred on day 2 (29°C, 1 Jun), day 4 (31°C, 3 Jun) and day 17 (31°C, 16 Jun, Fig S3). On each sampling day, one replicate fragment of each colony (n=10) and species (n=6) from each temperature treatment (n=2; n=120 total per sampling day) were used to measure net photosynthesis following the two-point method originally described and validated for *A. millepora* in Strahl et al. (2015). It is important to note that this method was not validated for additional species or under the experimental conditions used in the present study, nor were species specific photosynthesis-irradiance curves quantified to determine an appropriate saturating irradiance. Briefly, corals were incubated in enclosed 600-ml acrylic chambers at their respective treatment temperatures and light levels for 1.5 h. Chambers were placed onto custom built tables with rotating magnets, which served to power stir bars within each chamber to facilitate water mixing. For each run, four chambers without corals were used as blanks to account for potential changes in oxygen content due to the metabolic activity of other microorganisms in the seawater. For net photosynthesis measures, the O_2_ concentration of the seawater in each chamber was measured at the end of the run using a hand-held dissolved oxygen meter (HQ30d, equipped with LDO101 IntelliCAL oxygen probe, Hach, USA). Values from blank chambers were subtracted from measures made in coral chambers and the subsequent rate of net photosynthesis was related to coral surface area, calculated in µg O_2_/cm^2^/min.

To measure the fraction of autotrophically derived carbon translocated to host animals, five colony fragments of each species from each treatment (n=30 control, n=30 heat) were placed into 18-L of FSW with a 5-W aquarium pump for circulation and ^14^C-bicarbonate (specific activity: 56 mCi/mmol) was added to a final concentration of 0.28 µCi/ml. Corals were incubated for 5 hours in experimental tanks which experienced the normal experimental irradiance profile from 09:00 – 1400 GMT +10, rinsed with flow-through FSW for one hour to remove remaining unfixed ^14^C then snap frozen in liquid nitrogen.

Tissue was removed from snap frozen coral skeletons using an air gun and homogenized for 60 s using a Pro250 homogenizer (Perth Scientific Equipment, AUS). A 300-µl aliquot of the tissue homogenate was fixed with 5% formalin in FSW and used to quantify *Symbiodinium* cell density. The average cell number was obtained from four replicate haemocytometer counts of a 1-mm^3^ area and cell density was related to host protein content (as assessed below) and expressed as cells/mg host protein. Although surface area is common used as the normalization factor for *Symbiodinium* cell density, it has been recognized that areal abundance does not account for differences in host biomass (reviewed in (Cunning & Baker 2014). We found that normalization to soluble host protein more accurately reflected this difference in biomass, given that tissue thickness is significantly greater in *Goniopora columna* and not adequately accounted for by skeletal surface area normalization alone (Fig. S4). We determined the degree of bleaching, or the reduction in symbiont density in response to sustained thermal stress, by calculating the difference in symbiont cell densities between heat-treated and paired control samples following the full 17 days of experimental treatment and used this value as a proxy of holobiont fitness.

An additional 1-ml aliquot of holobiont homogenate was frozen at −20°C. The remaining homogenate was centrifuged for 2 min at 3500 rcf to separate host and symbiont fractions and 2-ml of the host tissue slurry was frozen at −20°C. Total soluble protein was quantified for host tissue samples in duplicate using a colorimetric assay (Bio-Rad Protein Assay Kit II, USA) following the manufacturer’s instructions. Coral skeletons were rinsed with 5% bleach then dried at room temperature. Skeletal surface area was quantified using the single wax dipping method (Veal et al. 2010) and skeletal volume (used to standardize respirometry chamber volumes) was determined by calculating water displacement in a graduated cylinder.

Sample radioactivity was determined using a liquid scintillation counter (Tri-Carb 2810TR v2.12, Perkin Elmer, USA). Triplicate host and duplicate holobiont 300 µl tissue homogenate aliquots were mixed with 3.5 ml Ultima Gold XR liquid scintillation cocktail (Perkin Elmer, USA). Samples were temperature and light adapted for 1 hour and then counted for 1.5 min using the default parameters. Counts per minute were converted to disintegrations per minute (DPM) using a standard curve derived from a ^14^C Ultima Gold Quench Standards Assay (Perkin Elmer, USA). Technical replicates were averaged. Host DPM values were divided by holobiont DPM values to yield the fraction of autotrophically derived carbon shared between partners, which we defined as our metric of host-symbiont cooperation. The degree of symbiont parasitism was calculated as the difference in this proportion of photosynthetically fixed carbon translocated to hosts between heat-treated and paired control samples following the full 17 days of experimental treatment, *sensu* Baker et al. (2018).

To identify major *Symbiodinium* clades hosted by focal species, DNA was extracted from symbiont fractions of the tissue homogenate for each replicate colony of each species (n=60, all control tank fragments) using Wayne’s method (Wilson et al. 2002). A restriction digest of the LSU region of *Symbiodinium* rRNA (Baker & Rowan 1997; Palstra 2000) consistently revealed single bands indicating the dominance of a single *Symbiodinium* clade for four of the six species (*A. millepora, M. aequituberculata, G. columna* and *P. lobata*) whereas communities in *G. astreata* and *G. acrhelia* appeared more variable (Fig. S5). Therefore, DNA was pooled in equal proportions by species (n=6 samples) to identify general species-specific communities using amplicon sequencing of the ITS2 region of *Symbiodinium* rRNA. Additional DNA samples for each of the *G. astreata* and *G. acrhelia* colony replicates (n=20 samples) were also submitted for sequencing at the Genome Sequencing and Analysis Facility at the University of Texas at Austin. Given the high diversity subsequently observed in pooled samples of *P. lobat*a and *M. aequituberculata*, additional colony replicate samples of each (n=3 and n=5, respectively) were later submitted for sequencing at Oregon State University’s Center for Genome Research and Biocomputing to compare symbiont community composition from individual samples.

### Amplicon Sequencing Analysis

For sample libraries prepared and sequenced at UT Austin’s GSAF, the ITS2 primers of Pochon et al. (Pochon et al. 2001) were used. For colony replicate samples sequenced at OSU’s CGRB (individual *P. lobata* and *M. aequituberculata* samples), the ITS2 primers of LaJeunesse (LaJeunesse 2002) were used to prepare libraries. All sequencing libraries were subsequently analyzed together. Prior to analysis, raw read data was filtered to remove reads which contained Illumina sequencing adapters or did not begin with the correct ITS2 amplicon primer sequence using the BBMAP package ver. 37.75 (http://sourceforge.net/projects/bbmap/). The DADA2 pipeline (Callahan 2016) implemented in R (Team 2017) was then used to infer sequence variants. Read data was analyzed according to the following tutorial (https://benjjneb.github.io/dada2/tutorial.html) with the following modifications: ITS2 primers were trimmed from the beginning of each read and forward and reverse reads were truncated at 210bp and 160bp respectively to remove low quality bases at the end of reads. Additional paired end reads were discarded if they exhibited more than one expected error or when a quality score of 2 or less was encountered. In addition, when inferring sequencing variants using the *dada*() command, the BAND_SIZE flag was set to 32 as is recommended for ITS data (https://benjjneb.github.io/dada2/tutorial.html). The distribution of raw read data per sample and reads lost during quality filtering and processing steps can be found in Table S1. Post-clustering curation of identified amplicon sequence variants (ASVs) was accomplished with the LULU algorithm, using the default parameters (Frøslev et al. 2017). The MCMC.OTU package (Green et al. 2014) was then used to remove sample outliers with low counts overall (z-score <-2.5) and remaining ASVs of abundance less than 0.001 (Quigley et al. 2014) prior to statistical analysis. In total, 12 ASVs were identified across samples that satisfied these criteria. These high confidence sequence variants were taxonomically classified through a blast search against the GeoSymbio ITS2 database https://sites.google.com/site/geosymbio/downloads (Franklin et al. 2012), and the best match was recorded. In cases where variants matched equally well to multiple references, all top hits were reported (Table S2). Resequencing of the individual *P. lobata* and *M. aequituberculata* samples indicated that the *P. lobata* pooled sample was contaminated at some stage of the sequencing process, as individual samples did not match the host pool (Fig. S6). In addition, two *Galaxea* samples (Gacr1 and Gast4) appeared mislabeled, based on the pattern of symbiont community differences among species (Fig S6) and we removed these samples prior to further analysis. To account for differences between individual samples and species pools, we calculated the normalized mean abundance of each variant across individual samples within a species and replaced the pooled sequence sample with this *in silico* pooled value, when available.

### Statistical Analyses

All statistical analyses were performed in R, version 3.4.2 (Team 2017). A series of linear mixed models were used to determine the effect of species, temperature treatment and sampling day on physiological metrics. We used a conservative approach to evaluate the effect of transmission mode on physiological metrics. Rather than modelling fixed effects of transmission mode directly, we modelled the fixed effects of species (levels: Amil, Maeq, Gcol, Plob, Gast, Gacr), sampling day (levels: day 2, day 4, day 17), temperature treatment (levels: heat, control) and all possible interactions on *Symbiodinium* cell density, net photosynthesis, and host-symbiont resource sharing using the *lme* command of the *nlme* package (Pinheiro et al. 2013), including source coral colony identity nested within reef of origin as a scalar random factor. All models were assessed for normality of residuals and homoscedasticity. Symbiont densities required a log-transformation to meet normality assumptions. To assess goodness-of-fit, we used the function *rsquared.glmm* to calculate the conditional R^2^ value for each of our mixed models (Johnson 2014). Significance of fixed factors within models was evaluated using Wald tests and Tukey’s post-hoc tests were used to evaluate significance among levels within factors and interactions when warranted. In these models, a significant effect of transmission mode would have been detected first as a significant difference among species, but significant differences in the hypothesized direction between each pair of vertical and horizontal transmitters in the subsequent Tukey’s tests were also required to satisfy reporting an effect of transmission mode overall.

To determine the impact of specific *Symbiodinium* community characteristics on responses among species, the DESeq package (Anders & Huber 2010) was used to construct a series of generalized linear models to evaluate differences in in the abundance of each ASV by species with respect to aspects of host-symbiont cooperation: mean initial symbiont density (Day 2, heat samples); the mean fraction of carbon shared by symbionts under ambient conditions (Day 2, 4, 17 control samples), the mean difference in the fraction of carbon shared by symbionts during bleaching (Day17, heat vs. control samples) and the mean difference in symbiont density during bleaching (Day17, heat vs. control samples). Models were run for 30 iterations and for models that did not converge, p-values were converted to NAs, the standard notation for missing data, prior to applying a multiple test correction (Benjamini & Hochberg 1995).

Predictive relationships between *Symbiodinium* cell density, the degree of bleaching, cooperation, and symbiont parasitism, in addition to *Symbiodinium* community diversity (quantified using the inverse Simpson index, (Magurran 2004)) and composition (the proportion of Clade D) were explored at the species level using a series of linear models. When a significant difference was detected among species for physiological trait comparisons, the model was re-run within each species and a multiple test correction using the method of Benjamini & Hochberg (1995) was applied. Analyses of symbiont community composition were run both using the full six species averages and excluding the species for which individual sample sequencing replicates were not available (*A. millepora* and *G. columna*). As models did not substantially differ, we report results for the full six species.

## RESULTS

### Physiological metrics of thermal tolerance and cooperation

Significant fixed effects of species, temperature treatment, sampling time and the time*treatment interaction were detected for the *Symbiodinium* cell density model (R^2^_GLMM_=0.53, Table S3). The fixed difference among species was driven by the low symbiont density on average in *P. lobata*, which differed significantly from densities in *A. millepora, M. aequituberculata, G. astreata* and *G. acrhelia* (Tukey’s HSD < 0.05, Fig. 1). No differences were detected in post-hoc tests between focal species pairs by symbiont transmission mode. Across species, cell densities did not differ between control and heat-treated corals on days 2 and 4, but were reduced in heat treated corals on day 17, by 5.4 x 10^5^ cells/mg host protein on average (Tukey’s HSD < 0.001, Fig. 1).

**Figure 1.**
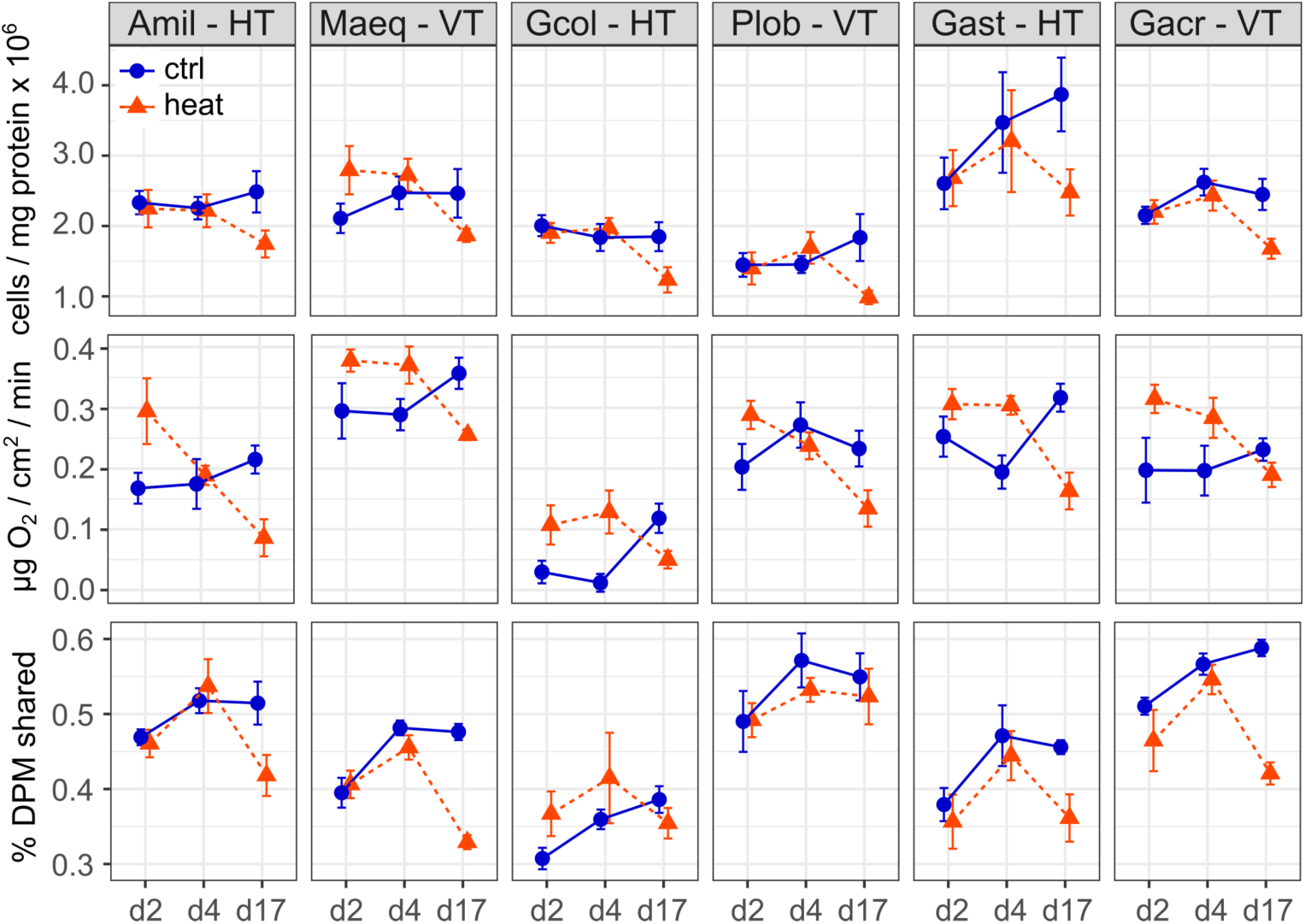
*Symbiodinium* cell density, expressed as cells per mg of host protein, rate of net photosynthesis, expressed as mg 0_2_ / cm^2^ / min and the proportion of photosynthetically fixed carbon translocated to the host, calculated as the ratio of total radioactivity in DPM in host to holobiont tissue fractions of focal coral species (Amil: *Acropora millepora*, Maqe: *Montipora aequituberculata*, Gcol: *Goniopora columna*, Plob: *Porites lobata*, Gast: *Galaxea astreata*, Gach: *Galaxea acrhelia*) under control (27°C, blue circles) and elevated (31°C, red triangles) temperature following 2, 4 and 17 days of treatment.

The model for changes in the rate of net photosynthesis identified significant effects of species, sampling time and the temperature treatment*time interaction (R^2^_GLMM_=0.53, Table S4). Average net O_2_ production rate was highest in *M. aequituberculata,* differing significantly from *A. millepora, G. columna, G. acrhelia,* and *P. lobata* Tukey’s HSD < 0.05, Fig. 1), and lowest in *G. columna*, significantly more so in comparison to *M. aequituberculata, P. lobata, G. astreata and G. acrhelia* (Tukey’s HSD < 0.001, Fig. 1). Net O_2_ production rate was also lower on average in *A. millepora* than in *M. aequituberculata* and *G. astreata* (Tukey’s HSD < 0.01, Fig. 1). Considering differences between focal species pairs by symbiont transmission mode, the rate of net photosynthesis was significantly higher in the vertically transmitting *M. aequituberculata* and *P. lobata* than in their horizontally transmitting counterparts, *A. millepora* and *G. columna*. However, it is important to note that absolute rates of oxygen production among focal species are much lower than previously reported values for corals (e.g. (Anthony et al. 2008). Corals were acclimated to a common light environment, but data on species-specific photosynthesis-irradiance curves are lacking. Consequently, the differences among species may be influenced by variation in species-specific photobiology.

Over the course of the experiment, net photosynthesis rates were elevated in heat-treated corals on days 2 and 4 relative to their respective controls (day2: 0.09 µg O_2_/cm^2^/min; day 4: 0.06 µg O_2_/cm^2^/min; Tukey’s HSD < 0.01, Fig. 1) and reduced in heat-treated corals on day 17, by 0.1 µg O_2_/cm^2^/min (Tukey’s HSD < 0.001, Fig. 1).

The proportion of carbon photosynthetically fixed by symbionts then translocated to hosts was significantly different among species, temperature treatment, sampling time, the treatment*time interaction and the species*treatment interaction (R^2^_GLMM_=0.73, Table S5). Carbon sharing in ambient conditions was significantly lower in *G. columna* in comparison to all other species (Tukey’s HSD < 0.05, Fig. 1) and highest in the vertically transmitting *G. acrhelia* and *P. lobata*, significantly more so than in *M. aequituberculata* and their horizontally transmitting counterparts, *G. astreata* and *G. columna* (Tukey’s HSD < 0.01, Fig. 1). Over time, no significant differences were detected in carbon sharing between control and heat-treated corals on days 2 and 4, but proportional translocation was significantly lower in heat-treated corals on day 17 (Tukey’s HSD < 0.001, Fig. 1). Relative to controls, carbon sharing decreased slightly under heat treatment on average in all species save *G. columna*, though the only significant difference was observed in *G. acrhelia* (heat vs. control Tukey’s HSD < 0.01, Fig. 1). Although the species*treatment*time interaction was not significant in the final model, this was likely driven by the decrease in heat-treated corals on day 17, which is most evident in *A. millepora, M. aequituberculata, G. astreata* and *G. acrhelia*.

### Relationships between symbiont density, degree of bleaching and carbon translocation

Symbiont cell density did not explain a significant proportion of the variance in carbon translocation under ambient conditions (control corals on days 2, 4 and 17) due to differences among species (F_1,5_ = 25.57, P < 0.001, Fig. 2). Within species, carbon translocation increased with increasing symbiont density in *G. acrhelia* (R^2^=0.22) but this relationship became non-significant after applying a multiple test correction.

**Figure 2.**
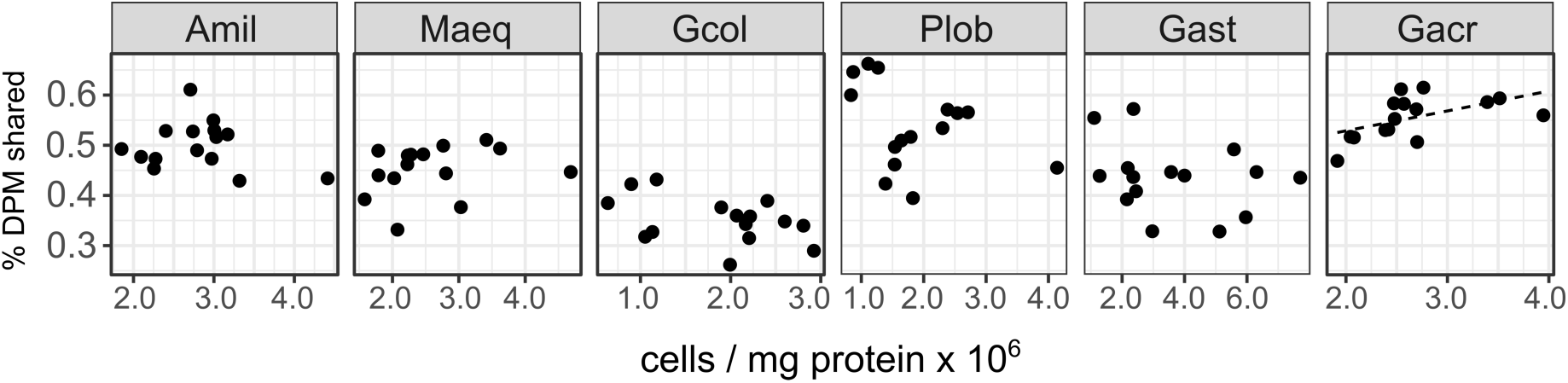
The proportion of photosynthetically fixed carbon translocated to the host in ambient conditions (27°C) as a function of *Symbiodinium* cell density by species (Amil: *Acropora millepora*, Maqe: *Montipora aequituberculata*, Gcol: *Goniopora columna*, Plob: *Porites lobata*, Gast: *Galaxea astreata*, Gach: *Galaxea acrhelia*).

Symbiont cell density in heat-treated corals on day 2 predicted a small portion of the variance in bleaching intensity on day 17 (calculated as the difference in cell density between paired control and heat-treated coral fragments). Greater bleaching was associated with higher initial symbiont cell densities (β = −0.34, R^2^ = 0.06, P = 0.04, Fig. 3A) and this relationship did not differ significantly among species. A similar, but non-significant trend was also observed between bleaching on day 17 and symbiont cell density on day 4 (β = −0.24, R^2^ = 0.04, P = 0.07).

**Figure 3.**
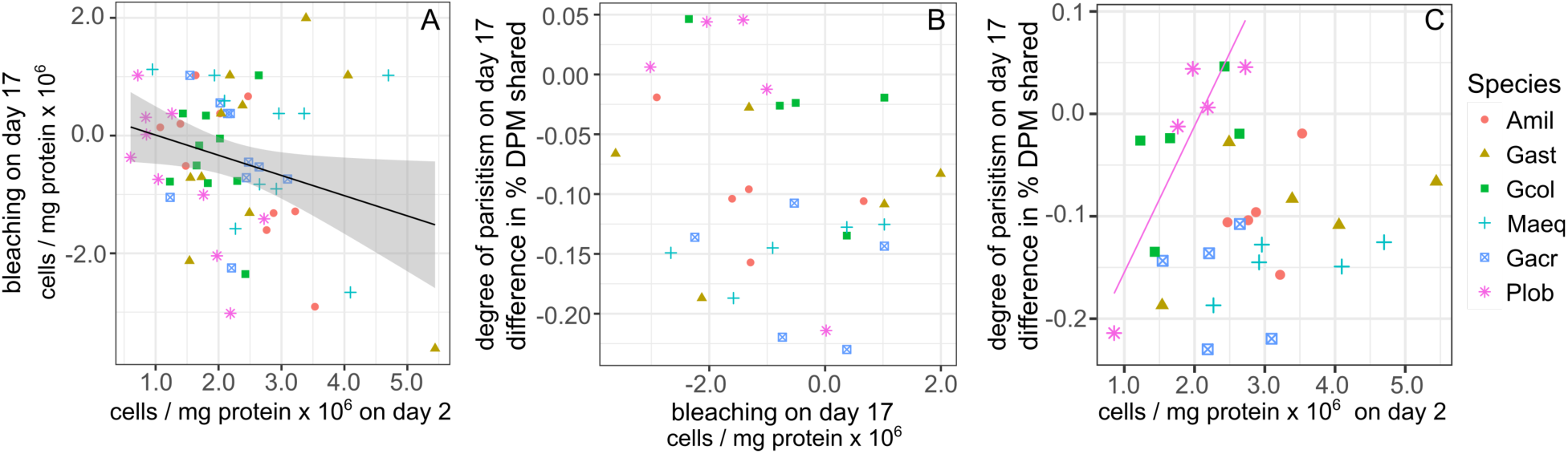
Relationships between changes in symbiont cell density and the proportion of photosynthetically fixed carbon translocated to hosts. (A) Bleaching intensity, calculated as the change in symbiont cell density between control (27°C) and heat-treated (31°C) samples on day 17 as a function of symbiont cell density in heat-treated corals on day 2. (B) The degree of symbiont parasitism, calculated as the difference in the proportion of photosynthetically fixed carbon translocated to hosts between control and heat-treated samples on day 17. (C) The degree of symbiont parasitism as a function of symbiont cell density in heat-treated corals on day 2. Regression line shown for Plob only. Amil: *Acropora millepora*, Maqe: *Montipora aequituberculata*, Gcol: *Goniopora columna*, Plob: *Porites lobata*, Gast: *Galaxea astreata*, Gach: *Galaxea acrhelia*.

No significant relationship was detected between the degree of bleaching and the degree of parasitism (calculated as the difference in carbon translocation between heat and control treated samples of paired corals by source colony) at the end of the experiment (Fig. 3B). Nor was the degree of parasitism explained by differences in symbiont cell density of heat-treated corals on day 2, but relationships differed among species (F_1,5_ = 6.88, P < 0.001). In *P. lobata*, a higher symbiont density on day 2 was associated with lower parasitism, or an increase in proportional carbon translocation in heat-treated corals on day 17, and this relationship remained marginally significant even after applying a multiple test correction (R^2^ = 0.89, P = 0.06, Fig 3C).

### The role of *Symbiodinium* community composition

Across all individual and pooled samples, 12 ASVs of sufficient representation (>0.1% abundance, *sensu* (Quigley et al. 2014) were identified and consisted of clade C and D-type *Symbiodinium* (Table S2, Fig. S6). *Symbiodinium* community composition varied among species (Fig. 4) but no relationships were detected between the abundance of individual ASVs and symbiont density, degree of bleaching or carbon translocation. Nor did we detect any relationship between *Symbiodinium* community diversity overall and symbiont density on day 2, mean carbon translocation under ambient conditions or the degree of bleaching at the end of the experiment (Fig. 5A,B,D). A marginally significant negative relationship was detected between community diversity and the degree of parasitism at the end of the experiment, where a more diverse community was associated with a greater degree of symbiont parasitism (R^2^ = 0.56, P = 0.054, Fig 5C). We did not detect any relationships between the percent of Clade D in the symbiont community and symbiont density on day 2, mean carbon translocation under ambient conditions, or the degree of parasitism or bleaching at the end of the experiment (Fig. 5E-H), but this was likely due to the fact that the abundance of D was very low in all species except *G. acrhelia*. *G. acrhelia,* however, did exhibit the highest carbon translocation rates under ambient conditions and the greatest transition towards parasitism on day 17 (Fig. 5F,G).

**Figure 4.**
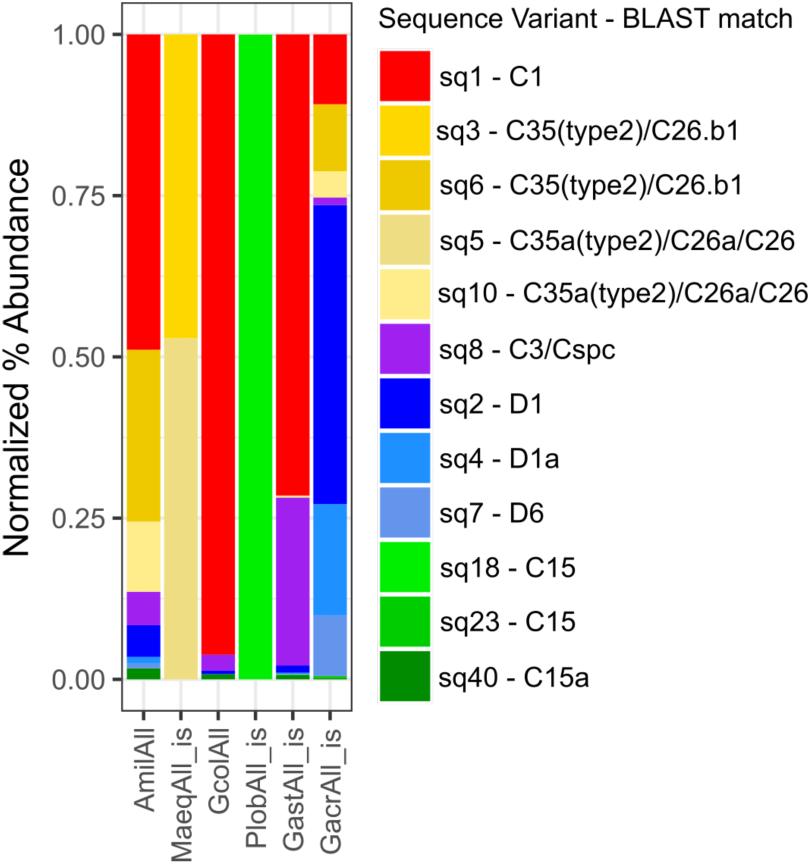
Normalized percent abundance of sequence variants by pooled sample. The ‘is’ label indicates samples for which sequence variant abundances across individually sequenced samples were pooled *in silico*. The best BLAST match against the GeoSymbio ITS2 database (Franklin et al. 2012) for each sequence variant is also reported.

**Figure 5.**
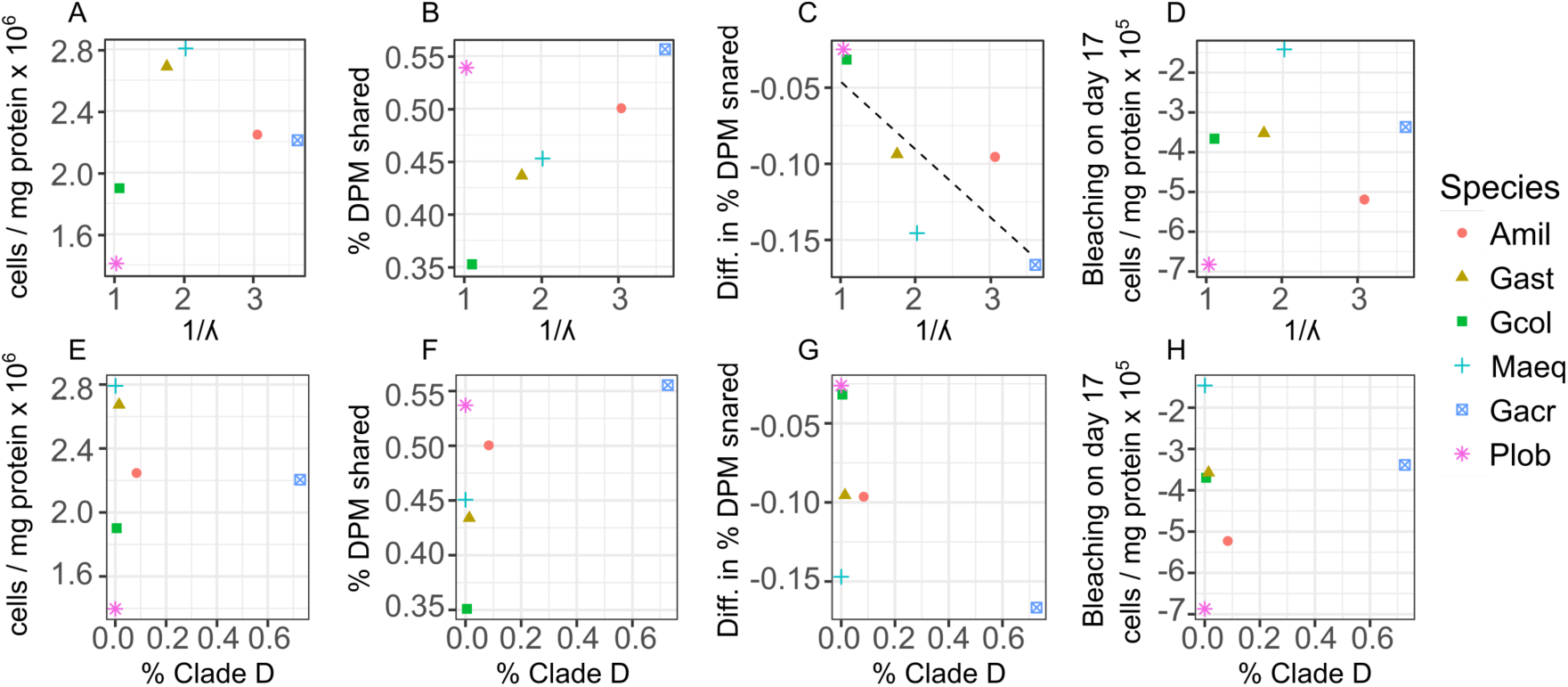
Relationships between changes in symbiont cell density or the proportion of photosynthetically fixed carbon translocated to hosts as a function of symbiont community diversity, expressed as the inverse Simpson index (A-D) or the normalized proportion of Clade D *Symbiodinium* (E-H) across species. For the dependent variables: (A,E) mean symbiont cell density in heat-treated corals (31°C) on day 2; (B,F) mean proportion of photosynthetically fixed carbon translocated to hosts under ambient conditions (27°C); (C,G) The degree of symbiont parasitism, calculated as the difference in the proportion of photosynthetically fixed carbon translocated to hosts between control and heat-treated corals on day 17.

## DISCUSSION

Understanding variation in the degree of cooperation between corals and their *Symbiodinium* will be critical for assessing survival potential among species and populations in the face of increasing environmental change (Lesser et al. 2013). As in other mutualisms (Ebert 2013; Frank 1994; Herre et al. 1999), vertical transmission has been proposed as an evolutionary mechanism for enhancing holobiont fitness in the Cnidarian-algal symbiosis (Putnam et al. 2012). However, we did not observe universally consistent differences in cooperation between vertical and horizontally transmitting species. The vertically transmitting *P. lobata* and *G. acrhelia* exhibited the highest levels of carbon translocation in ambient conditions, which we interpret as symbiont-host cooperation, significantly more so than their respective horizontally-transmitting counterparts, *G. columna* and *G. astreata*; but cooperation was not different between *M. aequituberculata* and *A. millepora*, and tended to be higher in the latter horizontally transmitting species. However, species-specific photosynthesis-irradiance curves were not measured and the potential for interactions between photophysiology and baseline rates of carbon translocation must also be explored. We also expected that the degree of breakdown in the host-symbiont relationship under heat stress, which we interpret as a transition towards parasitism *sensu* (Baker et al. 2018), would be comparatively intensified in horizontally transmitting species, but again, this was not the case. *P. lobata* did exhibit the least change in carbon translocation in spite of showing the same signs of bleaching as other species, but there was no difference in comparison to *G. columna* which also largely maintained its baseline translocation under elevated temperature. In addition, the greatest relative declines in the proportion of carbon translocated from symbionts to hosts actually occurred in the other two vertically transmitting species, *G. acrhelia* and *M. aequituberculata*.

Although we did not find significant support for the role of vertical transmission we still observed significant differences in cooperation among our six focal species, both baseline differences under ambient conditions and in the degree of transition towards parasitism under elevated temperature stress. We therefore conducted a series of *post hoc* analyses to explore other putative drivers of differential cooperation and thermal tolerance: differences in *Symbiodinium* cell density and/or in symbiont community composition.

The density of symbiont cells was recently proposed as a major driver underpinning the degree of cooperation between coral hosts and symbionts and the functional response of the coral symbiosis to environmental stressors (Cunning & Baker 2014). Studies in other species have shown that corals with greater initial *Symbiodinium* cell densities, as quantified by the symbiont to host cell ratio, are subsequently associated with greater bleaching severity in response to elevated thermal stress (Cunning & Baker 2013; Silverstein et al. 2014). This association has been hypothesized to result from the proportional increase in oxidative cellular stress: more symbionts yield more reactive oxygen species when the photosynthetic machinery is overloaded (Cunning & Baker 2014), though the recent work of Baker et al. (Baker et al. 2018), adds another potential explanation. They showed that under non-limiting nutrient conditions, *Symbiodinium* cell division in *Orbicella faveolata* actually increased in response to sub-bleaching temperature exposure, but that the metabolic costs were borne by the coral hosts (Baker et al. 2018), supporting the prediction that a transition to parasitism precedes unsustainable proliferation of the symbiont community which ultimately results in bleaching (Wooldridge 2009).

In examining relationships between symbiont cell densities, the intensity of the bleaching response at the end of our 17-day temperature exposure and carbon translocation rates, we did observe a negative relationship between initial symbiont cell density on days 2 and 4 and subsequent bleaching response on day 17, which did not significantly differ among our six focal species. However, initial symbiont cell densities did not predict the degree of the subsequent transition to parasitism. In fact, the majority of species showed a trend in which greater initial cell densities were associated with a greater maintenance of cooperation under bleaching stress. We also found no relationship between the degree of bleaching and the degree of parasitism on day 17, nor did symbiont cell density explain variation in cooperation among species in ambient conditions. For some species, cooperation tended to decrease with increasing density of symbionts, whereas in others it increased. Taken together, these observations support the association between initial symbiont cell density and subsequent bleaching intensity, but disagree with the proposed role of an alteration of host-symbiont cooperation in mediating the bleaching response.

The discrepancy in results among studies may be due to the importance of nutrient enrichment for observing a parasitic increase in symbiont communities, or in the difference in study duration and sampling design. Corals were exposed to a constant light environment, which likely did not provide a saturating irradiance across species. Consequently, alteration of carbon translocation as a function of species-specific photophysiology may explain baseline differences in ambient conditions, and future work should aim to test this hypothesis. In addition, while corals received supplemental feeding throughout the duration of this experiment, which likely introduced organic nitrogen, we did not explicitly manipulate inorganic nitrogen levels. It is also possible that variation among hosts in their ability to limit their symbionts’ nitrogen supply may have influenced the observed variation in the degree of parasitism (Cunning et al. 2017; Wooldridge 2009), though additional studies are needed to test this. In addition, Baker et al. (Baker et al. 2018) exposed corals to a +5°C temperature ramp over 8 days, sampling once 24 hours after the final temperature was reached, whereas the present study increased temperatures by +4°C over 5 days and maintained that difference for an additional 12 days, sampling at three time points, comparatively earlier and later. We did not observe an initial increase in symbiont cell density or decrease in carbon translocation on days 2 and 4 under elevated temperature, the experimental time-frame most analogous to that of Baker et al. (Baker et al. 2018). It is possible that these dynamics occurred during a time frame in which we did not sample; however, our final results argue against this explanation. On day 17, we observed symptoms of bleaching that did not differ across species: symbiont cell densities and rates of net photosynthesis were uniformly decreased. However, the transition to parasitism was not uniform, as some species exhibited significant differences in carbon translocation in response to heat stress whereas others did not. We therefore conclude that while symbiont density alone may be a reasonable predictor of the potential for observing a bleaching response under elevated temperature, it does not predict cooperative dynamics that likely also influence holobiont fitness.

We therefore also explored the role of symbiont community diversity on bleaching stress and cooperation. Predictions regarding the cooperative and fitness benefits of evolving vertical transmission are based on the assumption that the host-symbiont relationship becomes exclusive: symbiont population sizes are substantially reduced, resulting in genetic uniformity, more rapid co-evolution of partner traits and reduction in intra-symbiont community competition (Herre et al. 1999; Maynard Smith & Szathmary 1995). In this case, it may not be vertical transmission *per se* that influences host-symbiont cooperation, but the relative diversity of the symbiont community. A prior meta-analysis found that symbiont specificity was correlated with transmission mode, with horizontally transmitting species being more likely to interact with generalist symbionts (Fabina et al. 2012). However, the relationship between transmission mode and overall community diversity was not explored. Other more recent studies have also shown the potential for ontogenetic shifts in *Symbiodinium* community composition of putative vertical transmitters, potentially indicating the capacity for mixed-mode or cryptic horizontal transmission (Byler et al. 2013; Reich et al. 2017). Our results show that symbiont diversity does not partition by transmission mode. While communities in the vertically transmitting *M. aequituberculata* and *P. lobata* were largely uniform, consisting predominantly of C35 and C15-type sequence variants, respectively, *Symbiodinium* community diversity was more than twice as high in *G. acrhelia* in comparison to *G. astreata* (1/D = 3.6 vs. 1.7). In addition, community composition in the horizontally transmitting *G. columna* was also highly uniform, second only to that of *P. lobata* (1/D = 1.08 and 1.00, respectively).

In exploring the relationship between symbiont diversity at the ITS2 locus and metrics of host-symbiont cooperation and bleaching independent of transmission mode, we did not find any formally significant correlations, likely due to the fact that our symbiont community analysis was limited to the level of differences among the six species, greatly reducing our statistical power. However, there was a weak negative relationship between community diversity and the degree of parasitism under thermal stress. Species with the most genetically uniform symbiont communities, *P. lobata* and *G. columna*, maintained the highest levels of cooperation in spite of showing signs of bleaching. Yet there was no relationship between community diversity and baseline differences in cooperation among species under ambient conditions, as high rates of translocation were observed in species with both the lowest (*P. lobata*) and highest community diversity (*A. millepora, G. acrhelia*), though again, species-specific interactions between photophysiology and carbon translocation remain to be explored.

The presence of particular symbiont types has also been shown to influence holobiont fitness and carbon translocation. For example, conspecific corals hosting clade D *Symbiodinium* exhibit greater thermal tolerance than those hosting C1 or C2-types (Berkelmans & van Oppen 2006; Jones et al. 2008), however they generally grow more slowly under non-stressful conditions (Jones & Berkelmans 2010; Little et al. 2004) and receive less photosynthetically fixed carbon from their symbionts (Cantin et al. 2009). We found no significant relationships between the proportional abundance of Clade D-type symbionts and metrics of host-symbiont cooperation or bleaching. Most species had no, or a low proportion of Clade D, but the species with the greatest proportion of Clade D symbionts (*G. acrhelia*) exhibited both the highest carbon translocation under ambient conditions and the greatest transition to parasitism under elevated temperature stress. Similar to our observations regarding symbiont cell density, these results support prior observations that *Symbiodinium* community composition alone is not sufficient to explain variation in holobiont performance (Abrego et al. 2008; Baird et al. 2009a; Kenkel et al. 2013). However, we reiterate that our analyses are limited in comparing only averages among species. There was some variation observed in dominant symbiont types among individual coral colonies within species (Fig. S6) and a priority for future study should be to investigate whether these conclusions hold when considering intraspecific variation in symbiont community composition in addition to these broader interspecific differences.

Quantifying cooperation between symbiotic partners in terms of biologically realistic costs and benefits remains an outstanding question for many symbioses (Herre et al. 1999). The transfer of photosynthetically fixed carbon has long been known as a major cooperative benefit to the coral host as up to 95% of a coral’s energy requirements can be met through this mechanism (Muscatine 1990); however, reciprocal products shared by hosts with their symbionts remain largely unknown (Yellowlees et al. 2008). Similarly, heterotrophic feeding can offset the need for symbiont-derived carbon in some species and in these cases other symbiont-derived metabolites may be more critical for host fitness (Grottoli et al. 2006). Substantial variation in both intra- and inter-specific bleaching thresholds (Marshall & Baird 2000), suggests that levels of cooperation between host and symbiont may also vary. Significant work has gone into investigating coral bleaching over the past three decades, yet fundamental questions remain unresolved (Edmunds & Gates 2003). Ultimately, a greater understanding of both fine-scale interactions between coral hosts and symbionts and the evolutionary and ecological mechanisms that maintain and strengthen cooperation will be essential for managing these ecosystems (Davy et al. 2012; Lesser et al. 2013).

## CONCLUSIONS

This study investigated whether corals employing vertical symbiont transmission also exhibit enhanced cooperation and holobiont fitness. Contrary to theoretical predictions, we did not find significant support for the role of vertical transmission in spite of significant differences in cooperation among our six focal species. In a *post hoc* analysis of other drivers, we found that a greater initial symbiont cell density was associated with a greater bleaching intensity, but this association did not appear to result from an alteration of host-symbiont cooperation. Rather, the reduction in cooperation across species at the onset of bleaching was marginally associated with symbiont community diversity. The theoretical benefits of evolving vertical transmission are based on the underlying assumption that the host-symbiont relationship becomes genetically uniform, thereby reducing competition among symbionts. Taken together, our results suggest that it may not be vertical transmission *per se* that influences host-symbiont cooperation, but genetic uniformity of the symbiont community, though future work is needed to directly test this hypothesis.

## ACKNOWLEDGEMENTS

The authors gratefully acknowledge the efforts of A Bouriat in maintaining aquaria and processing coral samples. M Salmon, G Milton, A Severati, C Humphrey implemented the experimental aquaria design. Coral collection was accomplished with the help of S Noonan, V Mocellin, A Severati and M Nayfa. P Muir provided advice on coral taxonomic identification. Comments from J Caley, B Schaffelke and multiple anonymous reviewers greatly improved this manuscript.

## Supplementary Material

**Table S1.** Raw read counts and reads lost during filtering and clustering with DADA2.

| Sample | Raw | Input_DADA2 | filtered | denoised | merged | tabled | nonchim |
| --- | --- | --- | --- | --- | --- | --- | --- |
| AmilAll | 35820 | 35061 | 31194 | 31194 | 28315 | 28315 | 24013 |
| Gach1 | 74818 | 74054 | 67294 | 67294 | 66374 | 66301 | 62896 |
| Gach10 | 72354 | 71449 | 60322 | 60322 | 59915 | 59451 | 50573 |
| Gach2 | 74662 | 74012 | 63885 | 63885 | 62752 | 62250 | 50278 |
| Gach3 | 35176 | 34552 | 25626 | 25626 | 25323 | 25140 | 22602 |
| Gach4 | 93883 | 93234 | 80667 | 80667 | 79625 | 79064 | 63253 |
| Gach5 | 30755 | 30541 | 26595 | 26595 | 26375 | 26167 | 22976 |
| Gach6 | 40510 | 39898 | 34705 | 34705 | 34374 | 34019 | 31309 |
| Gach7 | 4472 | 4332 | 2643 | 2643 | 2611 | 2598 | 2528 |
| Gach8 | 33640 | 33267 | 29026 | 29026 | 28780 | 28443 | 26381 |
| Gach9 | 40512 | 40026 | 34308 | 34308 | 34026 | 33542 | 31669 |
| GachAll | 25286 | 24821 | 21705 | 21705 | 21399 | 21246 | 17958 |
| Gast1 | 1061 | 942 | 582 | 582 | 581 | 581 | 571 |
| Gast10 | 33882 | 33390 | 30573 | 30573 | 30231 | 30212 | 29162 |
| Gast2 | 21197 | 20863 | 18934 | 18934 | 18696 | 18696 | 17876 |
| Gast3 | 20029 | 19875 | 18266 | 18266 | 18003 | 18003 | 17431 |
| Gast4 | 16471 | 16248 | 13906 | 13906 | 13741 | 13558 | 12598 |
| Gast5 | 11943 | 11730 | 10196 | 10196 | 10172 | 10172 | 9182 |
| Gast6 | 45291 | 44813 | 40014 | 40014 | 39595 | 39574 | 38225 |
| Gast7 | 33652 | 33320 | 29961 | 29961 | 29883 | 29883 | 26478 |
| Gast8 | 23892 | 23539 | 21491 | 21491 | 21312 | 21287 | 20885 |
| Gast9 | 734 | 621 | 374 | 374 | 374 | 374 | 337 |
| GastAll | 31819 | 31304 | 27836 | 27836 | 27425 | 27367 | 23894 |
| GcolAll | 10196 | 9940 | 7972 | 7972 | 7839 | 7839 | 7661 |
| Maqe10 | 77568 | 69196 | 44998 | 44998 | 44705 | 44698 | 42679 |
| Maqe6 | 57788 | 46772 | 31275 | 31275 | 31160 | 31114 | 30225 |
| Maqe6b | 51170 | 27633 | 17137 | 17137 | 17074 | 17047 | 16564 |
| Maqe7 | 29361 | 23378 | 16091 | 16091 | 15970 | 15970 | 15338 |
| Maqe7b | 17131 | 16676 | 11653 | 11653 | 11567 | 11567 | 11491 |
| Maqe8 | 72073 | 43668 | 29519 | 29519 | 29346 | 29346 | 27746 |
| Maqe8b | 12976 | 12418 | 8388 | 8388 | 8314 | 8314 | 8214 |
| Maqe9 | 20282 | 19533 | 13318 | 13318 | 13226 | 13226 | 13131 |
| Maqe9b | 59314 | 31986 | 19854 | 19854 | 19715 | 19715 | 19002 |
| MaqeAll | 14445 | 14232 | 11940 | 11940 | 11754 | 11754 | 10322 |
| Plob6 | 16697 | 5454 | 3630 | 3630 | 725 | 584 | 561 |
| Plob6b | 28036 | 3924 | 2461 | 2461 | 1354 | 1294 | 1184 |
| Plob7 | 1469 | 454 | 263 | 263 | 44 | 30 | 3 |
| Plob7b | 17267 | 1798 | 1135 | 1135 | 197 | 109 | 109 |
| Plob8b | 36191 | 19086 | 12600 | 12600 | 11986 | 11940 | 11837 |
| PlobAll | 31619 | 30869 | 27527 | 27527 | 26407 | 26341 | 21754 |

**Table S2.** Best BLASTn matches to the GeoSymbio ITS2 database (Franklin et al. 2012) for curated amplicon sequence variants (ASVs) detected across samples. Value in % column indicates mean percent abundance across all samples in which the sequence variant, or sub-variants (identified through LULU curation), were identified. Match bit-scores are reported and hits separated by ‘/’ in GeoSymbio column indicate equally high scoring blast matches.

| ASV | % | Score | GeoSymbio BLAST |
| --- | --- | --- | --- |
| Sq1 | 31.6 | 523 | C1 |
| Sq2 | 18.8 | 501 | D1 |
| Sq3 | 10.6 | 508 | C35(type2)/C26.b1 |
| Sq4 | 7.0 | 501 | D1a |
| Sq5 | 11.4 | 508 | C35a(type2)/C26a/C26 |
| Sq6 | 6.6 | 512 | C35(type2)/C26.b1 |
| Sq7 | 3.8 | 501 | D6 |
| Sq8 | 4.9 | 523 | C3/Cspc |
| Sq10 | 2.5 | 512 | C35a(type2)/C26a/C26 |
| Sq18 | 1.6 | 520 | C15 |
| Sq23 | 0.8 | 523 | C15 |
| Sq40 | 0.3 | 523 | C15a |

**Table S3.** Wald test statistics for individual factors and interaction terms for the *Symbiodinium* cell density model. Significant terms are shown in bold.

|  | numDF | denDF | F-value | p-value |
| --- | --- | --- | --- | --- |
| --- | --- | --- | --- | --- |
| (Intercept) | 1 | 270 | 1.989512e+05 | 0.0000000 |
| **Species** | **5** | **52** | **9.116880e+00** | **0.0000028** |
| **Temp** | **1** | **270** | **1.260908e+01** | **0.0004528** |
| **Day** | **2** | **270** | **6.089511e+00** | **0.0025898** |
| Species:Temp | 5 | 270 | 1.104898e+00 | 0.3580558 |
| Species:Day | 10 | 270 | 1.452625e+00 | 0.1573259 |
| **Temp:Day** | **2** | **270** | **1.388694e+01** | **0.0000018** |
| Species:Temp:Day | 10 | 270 | 4.876186e-01 | 0.8975041 |

**Table S4.**
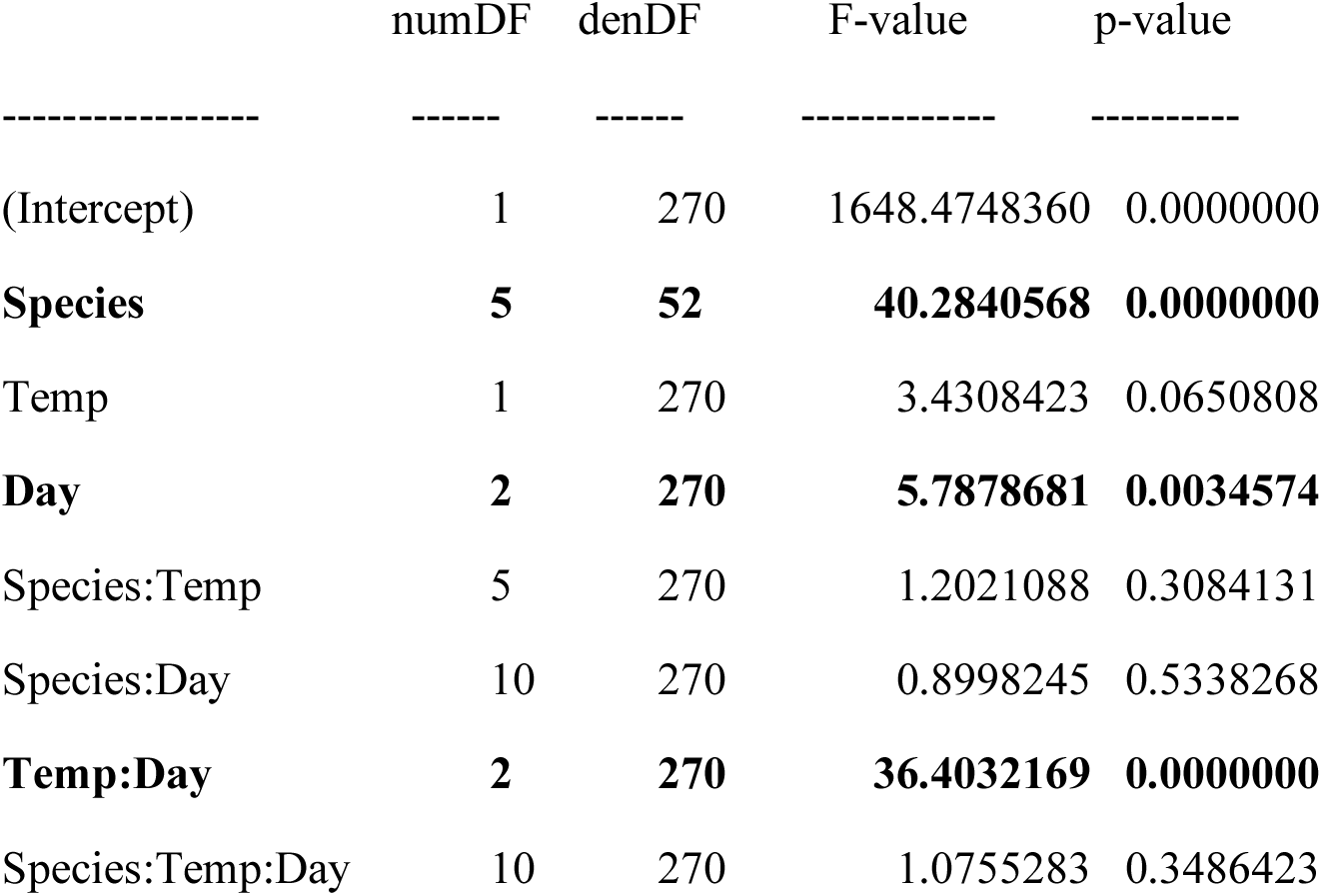
Wald test statistics for individual factors and interaction terms for the net photosynthesis model. Significant terms are shown in bold.

**Table S5.**
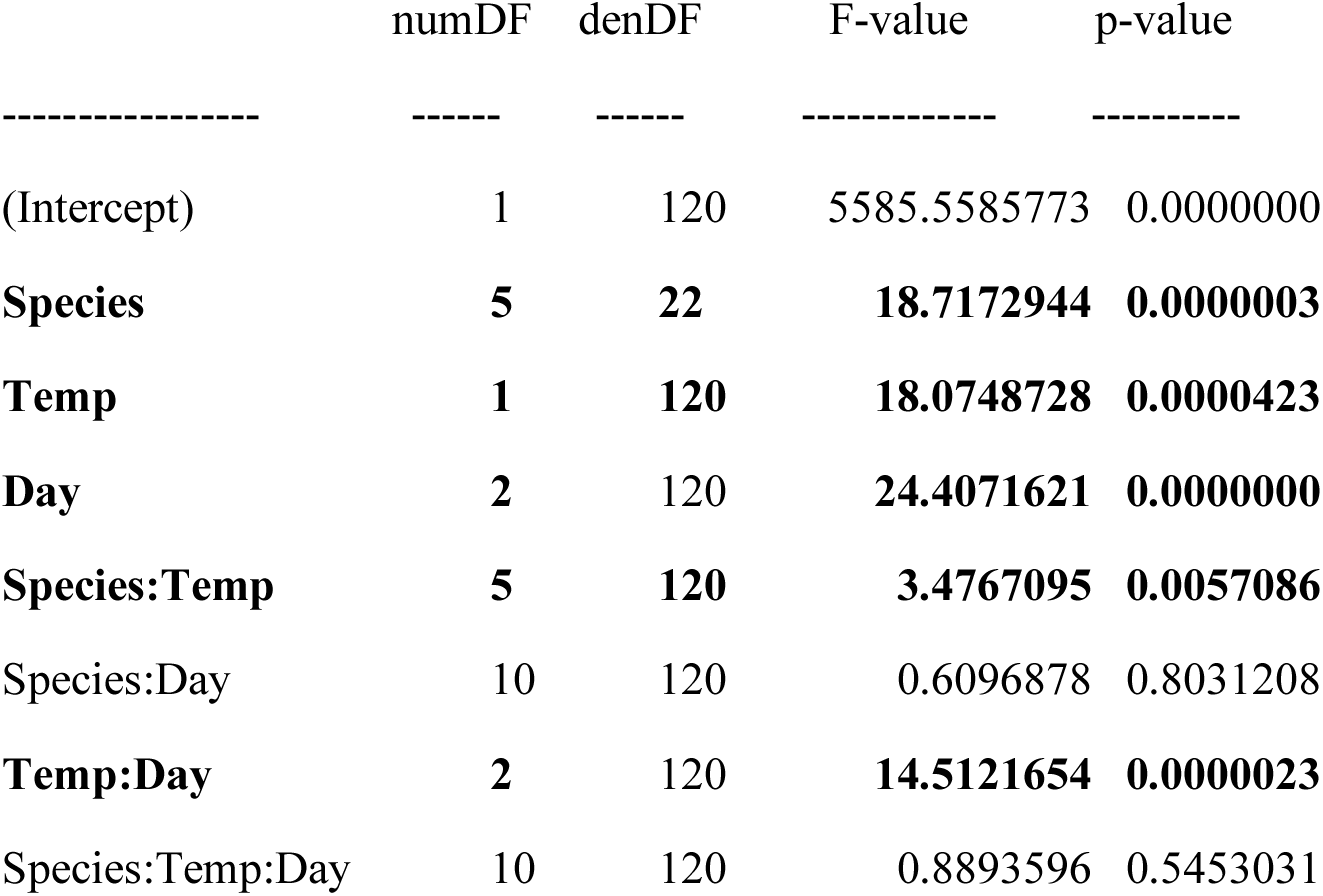
Wald test statistics for individual factors and interaction terms for the fraction of DPM shared model. Significant terms are shown in bold.

**Figure S1.**
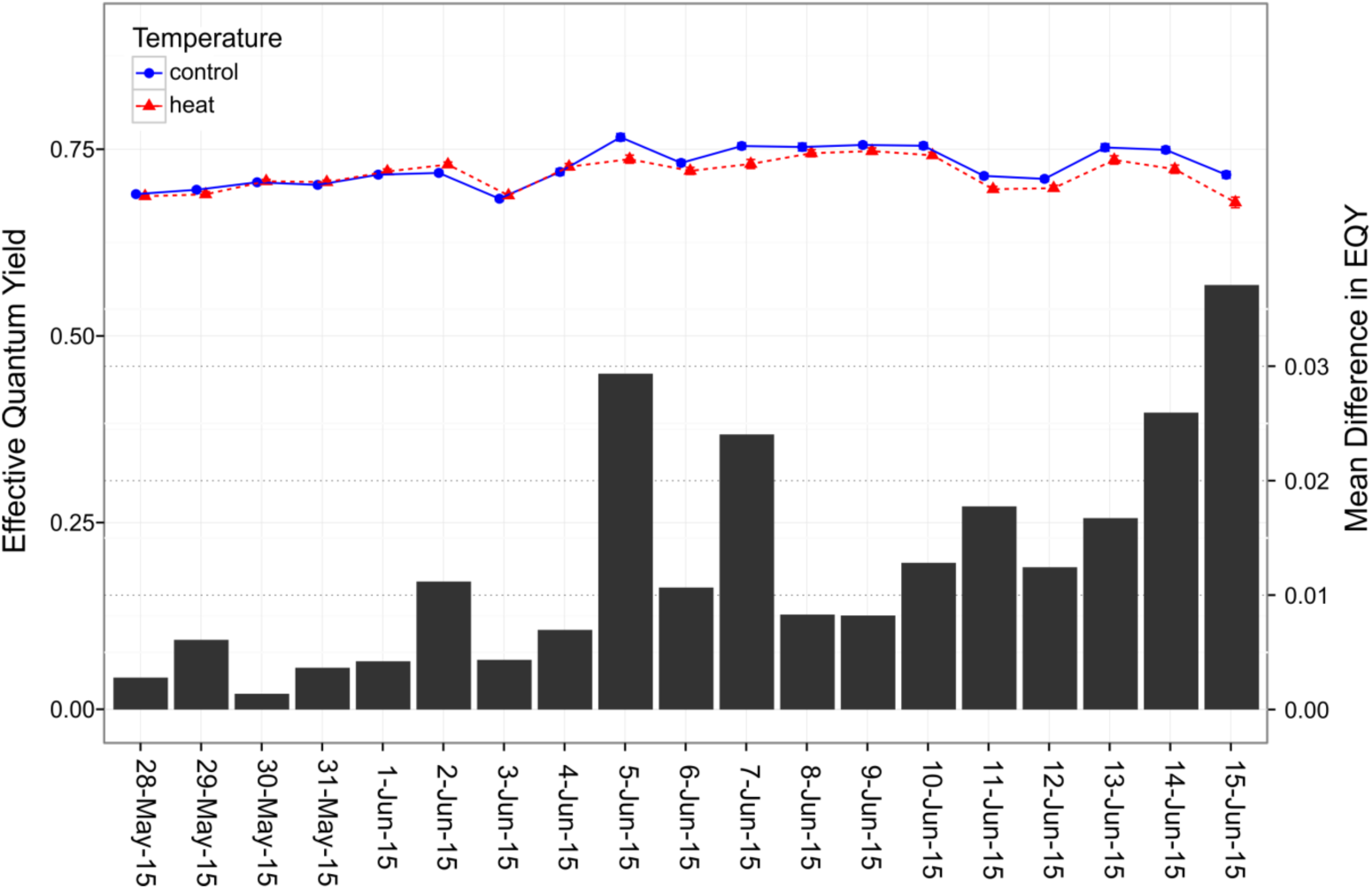
Daily measures of effective quantum yield (mean EQY±SEM, note that range of error bars is extremely small, in most cases barely exceeding points) of *Symbiodinium* photosystem II by temperature treatment. Bars show the difference in mean EQY between treatments through time (n=720 per bar).

**Figure S2.**
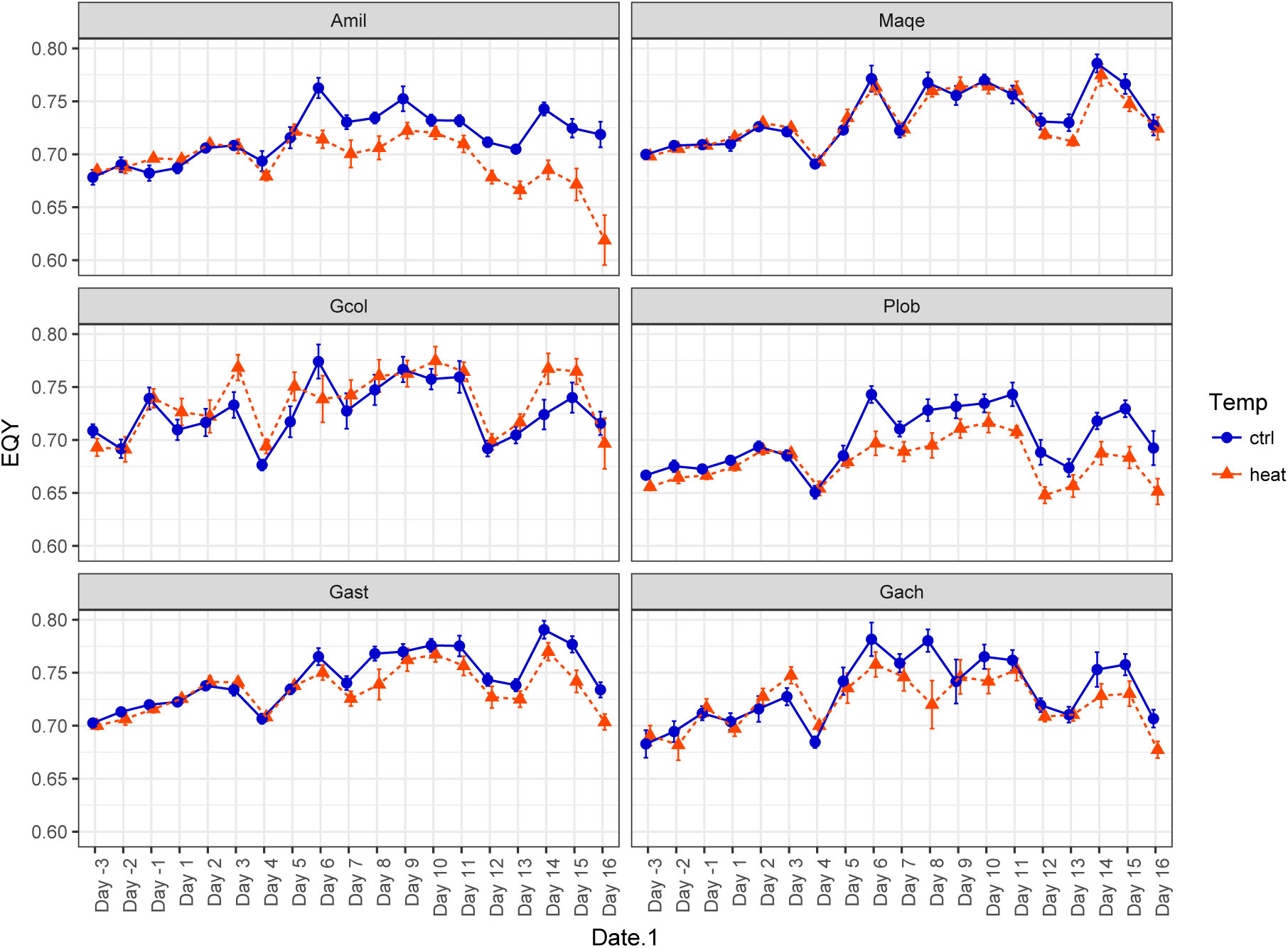
Daily measures of effective quantum yield (EQY±SEM) of *Symbiodinium* photosystem II by temperature treatment and species.

**Figure S3.**
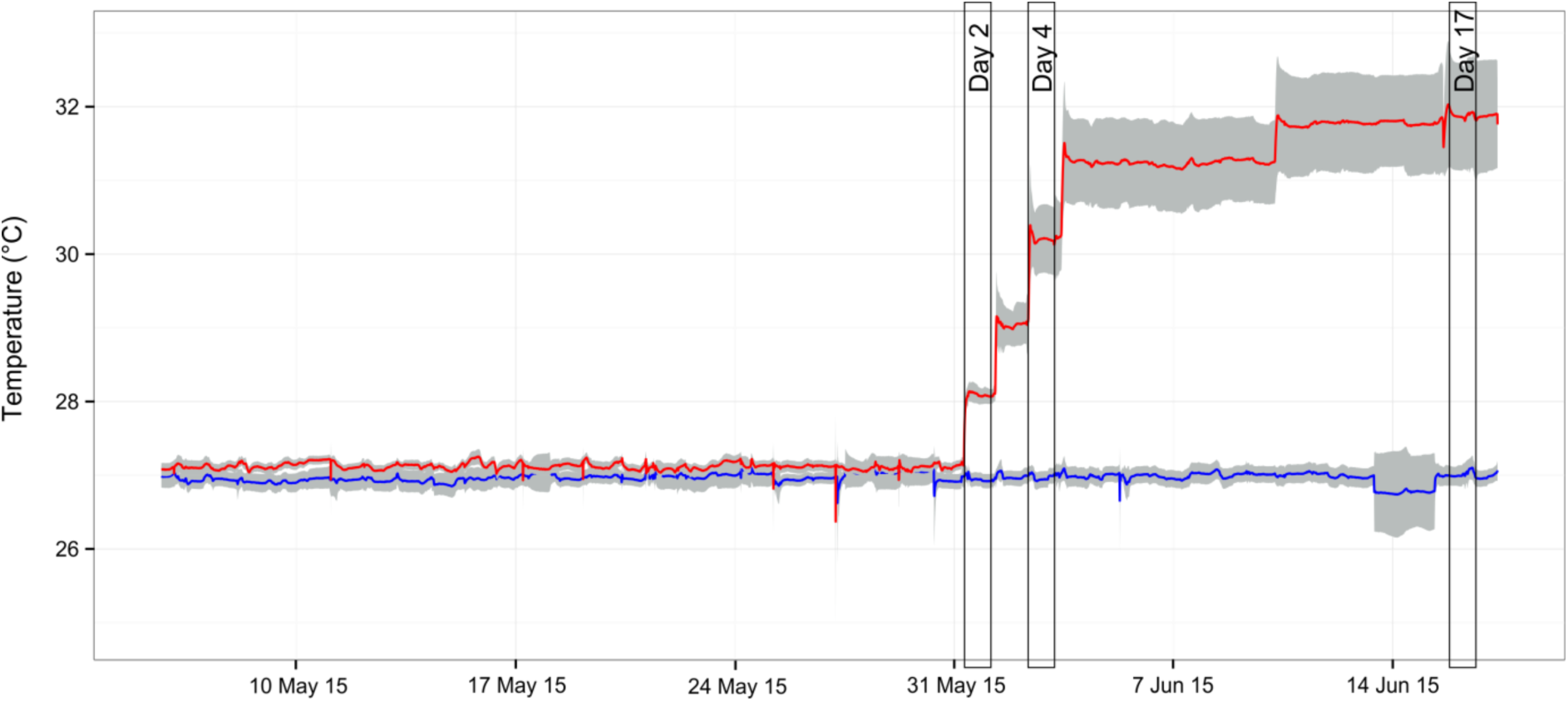
Temperature profile of experimental treatment tanks (±SD) by date. Bars indicate time of sampling.

**Figure S4.**
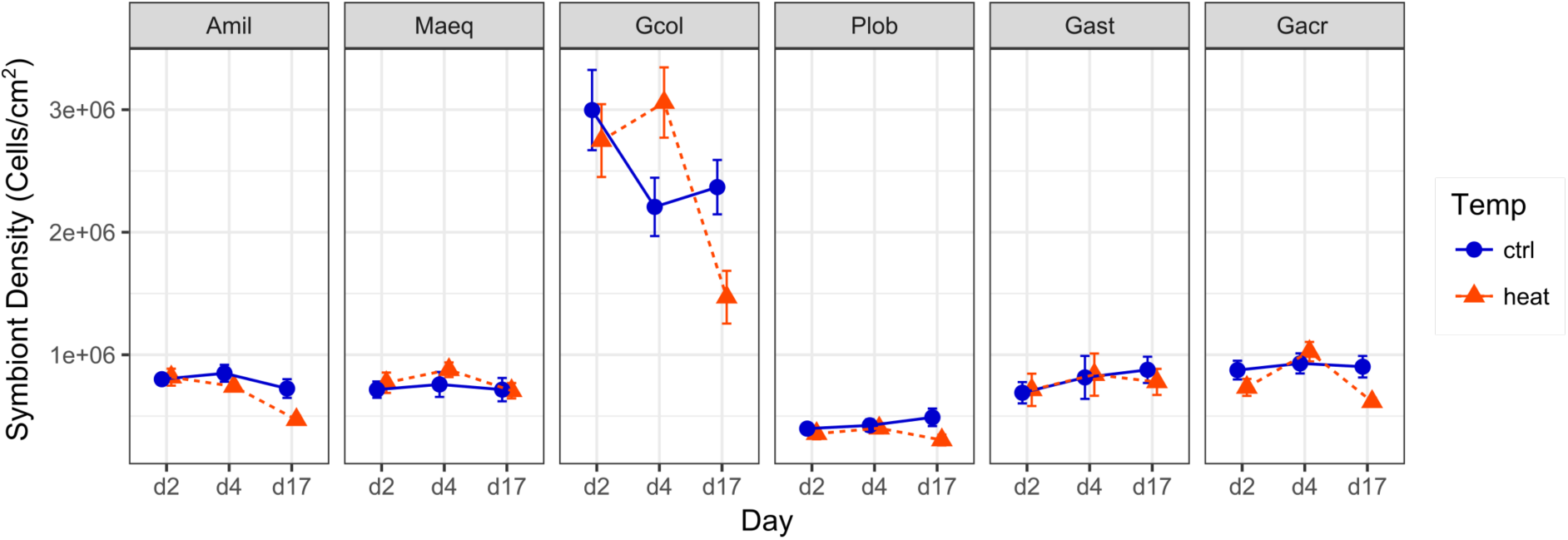
*Symbiodinium* cell density, expressed as cells per cm^2^ of coral surface area of focal coral species (Amil: *Acropora millepora*, Maeq: *Montipora aequituberculata*, Gcol: *Goniopora columna*, Plob: *Porites lobata*, Gast: *Galaxea astreata*, Gacr: *Galaxea acrhelia*) under control (27°C, blue circles) and elevated (31°C, red triangles) temperature following 2, 4 and 17 days of treatment.

**Figure S5.**
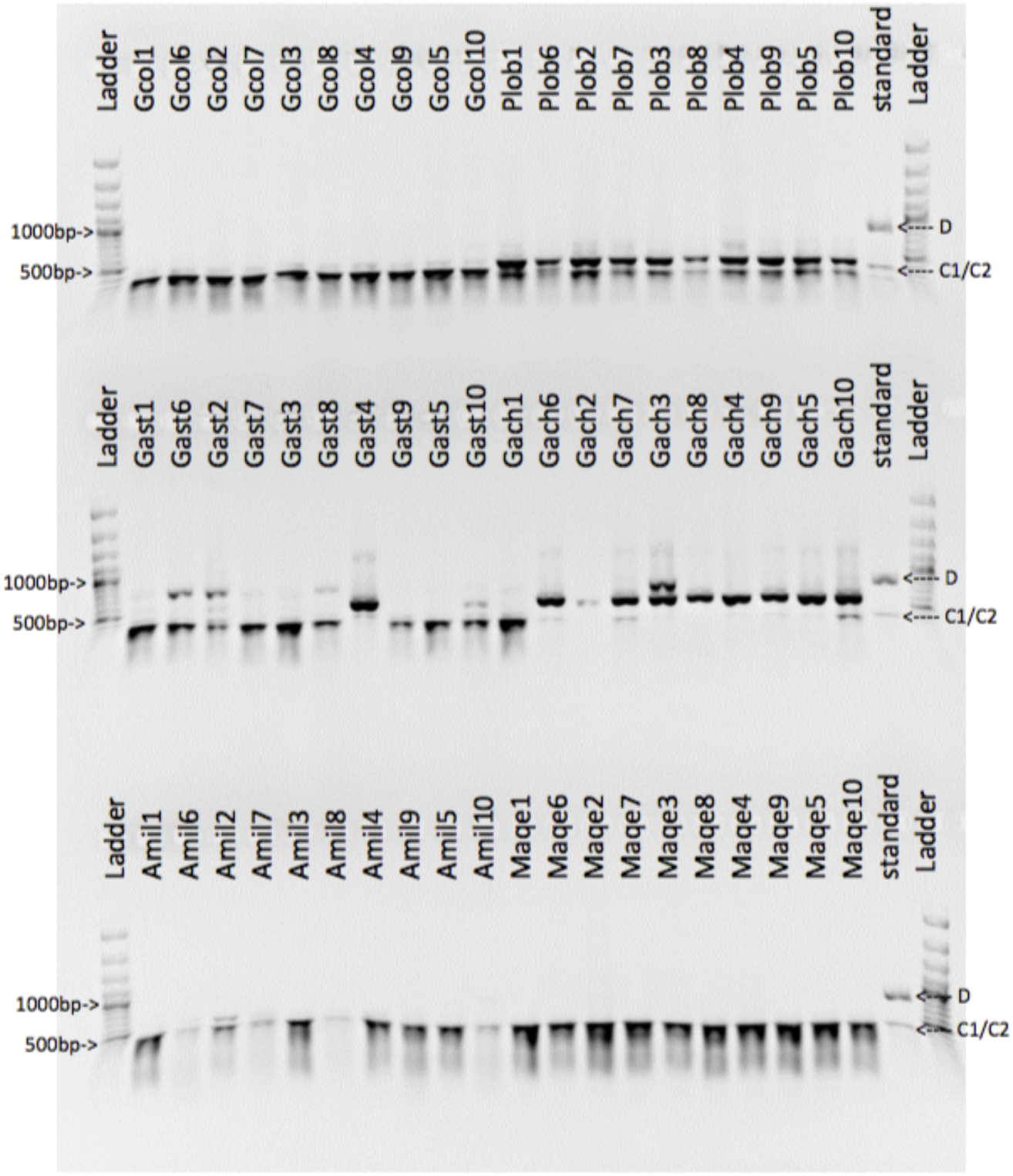
Taq1 digest of *Symbiodinium* LSU rRNA. Samples are grouped by species pairs. Standard shows expected banding pattern for *Symbiodinium* genotypes in clades C and D.

**Fig S6.**
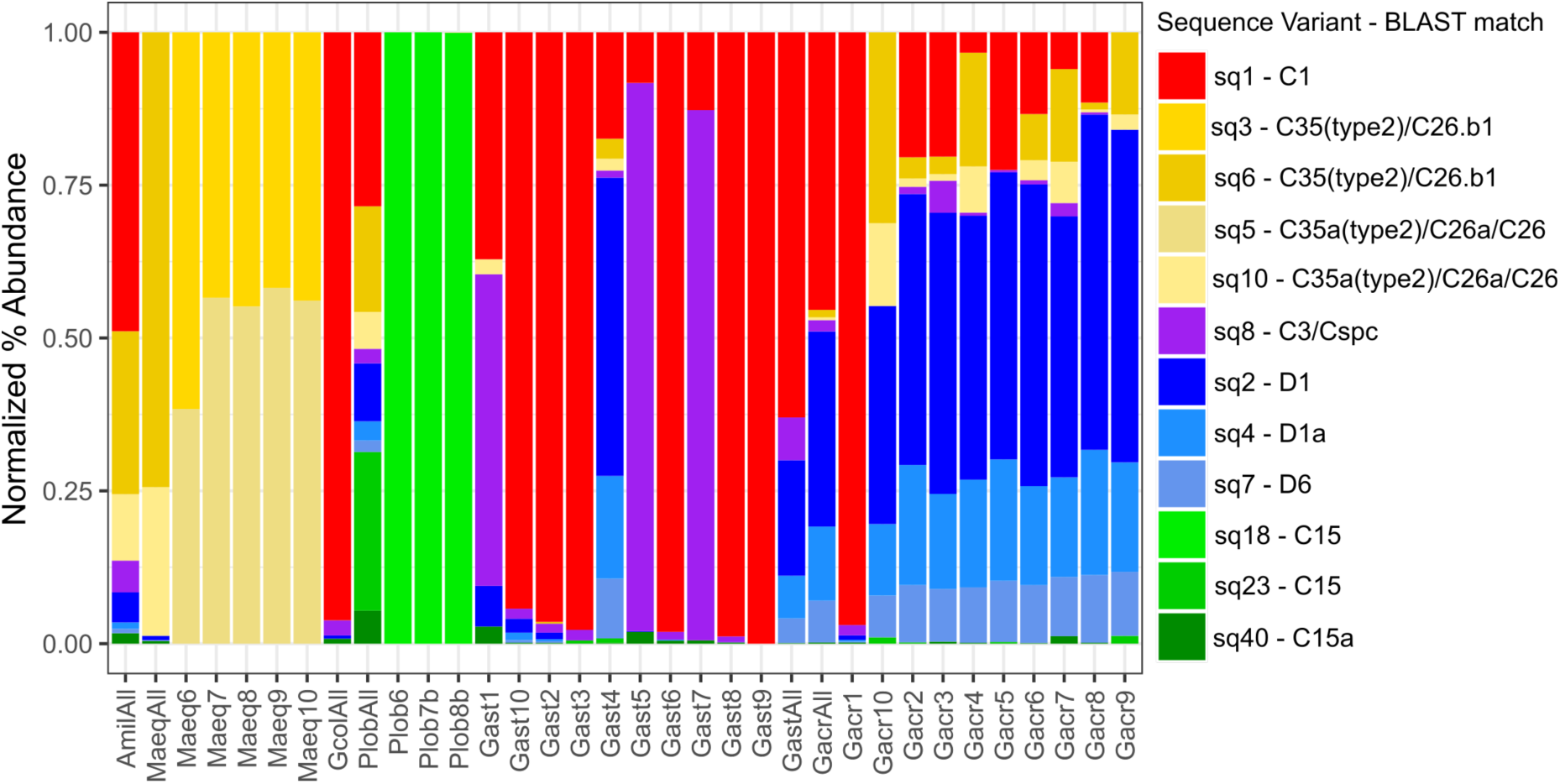
Normalized proportional abundance of sequence variants by samples sequenced at UT Austin’s GSAF (All and individual Gast/Gacr samples) and OSU’s CGRB (individual Maeq and Plob samples). Gacr1 and Gast4 were excluded from subsequent analyses as they appear to be mis-labeled and for analyses comparing differences among species, the ‘All’ samples for Gacr, Gast, Maeq and Plob were replaced by *in silico* averages calculated using the individual sample replicates shown here.

## REFERENCES

Abrego D, Ulstrup KE, Willis BL, and van Oppen MJH. 2008. Species-specific interactions between algal endosymbionts and coral hosts define their bleaching response to heat and light stress. Proceedings of the Royal Society B-Biological Sciences 275:2273–2282. 10.1098/rspb.2008.0180

Anders S, and Huber W. 2010. Differential expression analysis for sequence count data. Genome Biology 11:R106.

Anderson RM, and May RM. 1982. Coevolution of hosts and parasites. Parasitology 85:411–426.

Anthony KRN, Kline DI, Diaz-Pulido G, Dove S, and Hoegh-Guldberg O. 2008. Ocean acidification causes bleaching and productivity loss in coral reef builders. Proceedings of the National Academy of Sciences of the United States of America 105:17442–17446. 10.1073/pnas.0804478105

Baird AH, Bhagooli R, Ralph PJ, and Takahashi S. 2009a. Coral bleaching: the role of the host. Trends in Ecology & Evolution 24:16–20. 10.1016/j.tree.2008.09.005

Baird AH, Guest JR, and Willis BL. 2009b. Systematic and Biogeographical Patterns in the Reproductive Biology of Scleractinian Corals. Annual Review of Ecology Evolution and Systematics 40:551–571. 10.1146/annurev.ecolsys.110308.120220

Baker AC, and Rowan R. 1997. Diversity of symbiotic dinoflagellates (zooxanthellae) in scleractinian corals fo the Caribbean and Eastern Pacific. Proceedings of the 8th International Coral Reef Symposium. p 1301–1306.

Baker DM, Freeman CJ, Wong JCY, Fogel ML, and Knowlton N. 2018. Climate change promotes parisitism in a coral symbiosis. The ISME Journal. https://doi.org/10.1038/s41396-018-0046-8

Benjamini Y, and Hochberg Y. 1995. Controlling the false discovery rate - a practical and powerful approach to multiple testing. Journal of the Royal Statistical Society Series B-Methodological 57:289–300.

Berkelmans R, and van Oppen MJH. 2006. The role of zooxanthellae in the thermal tolerance of corals: a ‘nugget of hope’ for coral reefs in an era of climate change. Proceedings of the Royal Society B-Biological Sciences 273:2305–2312. 10.1098/rspb.2006.3567

Bright M, and Bulgheresi S. 2010. A complex journey: transmission of microbial symbionts. Nature Reviews Microbiology 8:218–230.

Brown BE. 1997. Coral bleaching: causes and consequences. Coral Reefs 16:S129–S138.

Bull JJ. 1994. Perspective: virulence. Evolution 48:1423–1437.

Bull JJ, Molineux IJ, and Rice WR. 1991. Selection of benevolence in a host-parasite system. Evolution 45:857–882.

Byler KA, Carmi-Veal M, Fine M, and Goulet TL. 2013. Multiple symbiont acquisition strategies as an adaptive mechanism in the coral Stylophora pistillata. PLoS ONE 8:e59596.

Callahan BJ, McMurdie, Paul J, Rosen, Michael J, Han, Andrew W, Johnson, Amy Jo A, Holmes, Susan P. 2016. DADA2: High-resolution sample inference from Illumina amplicon data. Nature Methods 13:581–583.

Cantin NE, van Oppen MJH, Willis BL, Mieog JC, and Negri AP. 2009. Juvenile corals can acquire more carbon from high-performance algal symbionts. Coral Reefs 28:405–414. 10.1007/s00338-009-0478-8

Cunning R, and Baker AC. 2013. Excess of algal symbionts increase the susceptibility of reef corals to bleaching. Nature Climate Change 3:259–262.

Cunning R, and Baker AC. 2014. Not just who, but how many: the importance of partner abundance in reef coral symbioses. Frontiers in Microbiology 5. 10.3389/fmicb.2014.00400

Cunning R, Muller EB, Gates RD, and Nisbet RM. 2017. A dynamic bioenergetic model for coral-Symbiodinium symbioses and coral bleaching as an alternate stable state. Journal of Theoretical Biology 431:49–62. 10.1016/j.jtbi.2017.08.003

Daly M, Brugler MR, Cartwright P, Collines AG, Dawson MN, Fautin DG, France SC, McFadden CS, Opresko DM, Rodriguez E, Romano SL, and Stake JL. 2007. The phylum Cnidaria: A review of phylogenetic patterns and diversity 300 years after Linnaeus. In: Zhang ZQ, and Shear WA, eds. Linnaeus Tercentenary: Progress in Invertebrate Taxonomy: Zootaxa, 127–182.

Davy SK, Allemand D, and Weis VM. 2012. Cell biology of Cnidarian-dinoflagellate symbiosis. Microbiology and Molecular Biology Reviews 76:229–261.

Doebeli M, and Knowlton N. 1998. The evolution of interspecific mutualisms. Proceedings of the National Academy of Sciences of the United States of America 95:8676–8680.

Dusi E, Gougat-Barbera C, Berendonk TU, and Kaltz O. 2015. Long-term selection experiment produces breakdown of horizontal transmissibility in parasite with mixd transmission mode. Evolution 69:1069–1076.

Ebert D. 2013. The epidemiology and evolution of symbionts with mixed-mode transmission. Annual Review of Ecology, Evolution, and Systematics 44:7.1–7.21.

Ebert D, and Bull JJ. 2003. Challenging the trade-off model for the evolution of virulence: is virulence management feasible? Trends in Microbiology 11:15–20.

Edmunds PJ, and Gates RD. 2003. Has coral bleaching delayed our understanding of fundamental aspects of coral-dinoflagellate symbioses? BioScience 53:976–980.

Fabina NS, Putnam HM, Franklin EC, Stat M, and Gates RD. 2012. Transmission mode predicts specificity and interaction patterns in coral-Symbiodinium networks. PLoS ONE 7:e44970. doi10.1371/journal.pone.0044970

Frank SA. 1994. Genetics of mutualism: the evolution of altruism between species. Journal of Theoretical Biology 170:393–400.

Franklin EC, Stat M, Pochon X, Putnam HM, and Gates RD. 2012. GeoSymbio: a hybrid, cloud-based web application of global geospatial bioinformatics and ecoinformatics for Symbiodinium-host symbioses. Molecular Ecology Resources 12:369–373.

Friesen ML, and Jones EI. 2013. Modelling the evolution of mutualistic symbioses. In: van Helden J, Toussaint A, and Thieffry D, eds. Bacterial Molecular Networks: Methods and Protocols: Springer Science+Business Media, 481–499.

Frøslev TG, Kjøller R, Bruun HH, Ejrnæs R, Brunbjerg AK, Pietroni C, and Hansen AJ. 2017. Algorithm for post-clustering curation of DNA amplicon data yields reliable biodiversity estimates. Nature Communications 8:1188. 10.1038/s41467-017-01312-x

Green EA, Davies SW, Matz MV, and Medina M. 2014. Quantifying cryptic Symbiodinium diversity with Orbicella faveolata and Orbicella franksi at the Flower Garden Banks, Gulf of Mexico. PeerJ 2:e386. 10.7717/peerj.386

Grottoli AG, Rodrigues LJ, and Palardy JE. 2006. Heterotrophic plasticity and resilience in bleached corals. Nature 440:1186–1189. 10.1038/nature04565

Harrison PL, and Wallace CC. 1990. Reproduction, dispersal and recruitment of scleractinian corals. In: Dubinsky Z, ed. Ecosystems of the World: Coral Reefs. New York: Elsevier, 133–207.

Hartmann AC, Baird AH, Knowlton N, and Huang D. 2017. The paradox of environmental symbiont acquisition in obligate mutualisms. Current Biology 27:3711–3716.

Hatcher BG. 1988. Coral reef primary productivity: A beggar’s banquet. Trends in Ecology and Evolution 3:106–111.

Herre EA. 1993. Population structure and the evolution of virulence in nematode parasites of fig wasps. Science 259:1442–1445.

Herre EA, Knowlton N, Mueller UG, and Rehner SA. 1999. The evolution of mutualisms: exploring the paths between conflict and cooperation. Trends in Ecology & Evolution 14.

Hoegh-Guldberg O, Mumby PJ, Hooten AJ, Steneck RS, Greenfield P, Gomez E, Harvell CD, Sale PF, Edwards AJ, Caldeira K, Knowlton N, Eakin CM, Iglesias-Prieto R, Muthiga N, Bradbury RH, Dubi A, and Hatziolos ME. 2007. Coral reefs under rapid climate change and ocean acidification. Science 318:1737–1742. 10.1126/science.1152509

Hughes TP, Baird AH, Bellwood DR, Card M, Connolly SR, Folke C, Grosberg R, Hoegh-Guldberg O, Jackson JBC, Kleypas J, Lough JM, Marshall P, Nystrom M, Palumbi SR, Pandolfi JM, Rosen B, and Roughgarden J. 2003. Climate change, human impacts, and the resilience of coral reefs. Science 301:929–933.

Johnson PCD. 2014. Extension of Nakagawa & Schielzeth’s R2GLMM to random slopes models. Methods in Ecology and Evolution 5:944–946.

Jones A, and Berkelmans R. 2010. Potential Costs of Acclimatization to a Warmer Climate: Growth of a Reef Coral with Heat Tolerant vs. Sensitive Symbiont Types. PLoS ONE 5. 10.1371/journal.pone.0010437

Jones A, Berkelmans R, van Oppen MJH, Mieog JC, and Sinclair W. 2008. A community change in the algal endosymbionts of a scleractinian coral following a natural bleaching event: field evidence of acclimatization. Proceedings of the Royal Society B-Biological Sciences 275:1359– 1365.

Kenkel C, Goodbody-Gringley G, Caillaud D, Davies SW, Bartels E, and Matz M. 2013. Evidence for a host role in thermotolerance divergence between populations of the mustard hill coral (Porites astreoides) from different reef environments. Molecular Ecology 22:4335–4348.

Kerr AM, Baird AH, and Hughes TP. 2011. Correlated evolution of sex and reproductive mode in corals (Anthozoa: Scleractinia). Proceedings of the Royal Society B-Biological Sciences 278:75–81.

Kiers TE, and West SA. 2015. Evolving new organisms via symbiosis. Science 348:392–394.

Knowlton N, Brainard RE, Fisher R, Moews M, Plaisance L, and Caley MJ. 2010. Coral Reef Biodiversity. In: McIntyre A, ed. Life in the World’s Oceans: Diversity, Distribution, and Abundance. Oxford: Blackwell Publishing Ltd., 65–78.

LaJeunesse TC. 2002. Diversity and community structure of symbiotic dinoflagellates from Caribbean coral reefs. Marine Biology 141:387–400.

Lesser MP, Stat M, and Gates RD. 2013. The endosymbiotic dinoflagellates (Symbiodinium sp.) of corals are parasites and mutualists. Coral Reefs 32:603–611.

Little AF, van Oppen MJH, and Willis BL. 2004. Flexibility in algal endosymbioses shapes growth in reef corals. Science 304:1492–1494.

Magurran AE. 2004. Measuring biological diversity. Oxford, UK: Blackwell Science Ltd.

Marshall PA, and Baird AH. 2000. Bleaching of corals on the Great Barrier Reef: differential susceptibilities among taxa. Coral Reefs 19:155–163.

Maynard Smith J, and Szathmary E. 1995. *The Major Transitions in Evolution*. Oxford: Freeman.

Moran NA. 2006. Symbiosis. Current Biology 16:R866–R871.

Muscatine L. 1990. The role of symbiotic algae in carbon and energy flux in reef corals. In: Dubinsky, ed. Ecosystems of the world 25: Coral reefs. New York: Elsiever, 75–87.

Nowak MA, Bonhoeffer S, and May RM. 1994. Spatial games and the maintenance of cooperation. Proceedings of the National Academy of Sciences of the United States of America 91:4877– 4881.

Palstra F. 2000. Host-endosymbiont specificity in Acropora corals of the Indo-Pacific? James Cook University.

Pinheiro J, Bates D, DebRoy S, Sarkar D, and Team RDC. 2013. nlme: Linear and nonlinear mixed effects models. R package version 3.1-113 ed.

Pochon X, Pawlowski J, Zaninetti L, and Rowan R. 2001. High genetic diversity and relative specificity among Symbiodinium-like endosymbiotic dinoflagellates in soritid foraminiferans. Marine Biology 139:1069–1078.

Putnam HM, Stat M, Pochon X, and Gates RD. 2012. Endosymbiotic flexibility associates with environmental sensitivity in scleractinian corals. Proceedings of the Royal Society B: Biological Sciences 279:4352–4361.

Quigley KM, Davies SW, Kenkel CD, Willis BL, Matz MV, and Bay LK. 2014. Deep-Sequencing Method for Quantifying Background Abundances of Symbiodinium Types: Exploring the Rare Symbiodinium Biosphere in Reef-Building Corals. PLoS ONE 9:e94297. doi:10.1371/journal.pone.0094297

Ralph PJ, Larkum AWD, and Kuhl M. 2005. Temporal patterns in effective quantum yield of individual zooxanthellae expelled during bleaching. Journal of Experimental Marine Biology and Ecology 316:17–28.

Reich HG, Robertson DL, and Goodbody-Gringley G. 2017. Do the shuffle: Changes in Symbiodinium consortia throughout juvenile coral development. PLoS ONE 12:e0171768. 10.1371/journal.pone.0171768

Roth MS. 2014. The engine of the reef: photobiology of the coral-algal symbiosis. Frontiers in Microbiology 5:422. 10.3389/fmicb.2014.00422

Sachs JL, Essenberg CJ, and Turcotte MM. 2011. New paradigms for the evolution of beneficial infections. Trends in Ecology & Evolution 26:202–209.

Sachs JL, Mueller UG, Wilcox TP, and Bull JJ. 2004. The evolution of cooperation. Quarterly Review of Biology 79:135–160.

Sachs JL, and Wilcox TP. 2006. A shift to parasitism in the jellyfish symbiont Symbiodinium microadriaticum. Proceedings of the Royal Society B-Biological Sciences 273:425–429.

Silverstein RN, Cunning R, and Baker AC. 2014. Change in algal symbiont communities after bleaching, not prior heat exposure, increases heat tolerance of reef corals. Global Change Biology 21:236–249.

Stanley GD, and Swart PK. 1995. Evolution of the coral-zooxanthellae symbiosis during the Triassic: a geochemical approach. Paleobiology 21:179–199.

Stewart AD, Logsdon JM, and Kelley SE. 2005. An empirical study of the evolution of virulece under both horizontal and vertical transmission. Evolution 59:730–739.

Strahl J, Stolz I, Uthicke S, Vogen N, Noonan SHC, and Fabricius KE. 2015. Physiological and ecological performance differs in four coral taxa at a volcanic carbon dioxide seep. Comparative Biochemistry and Physiology Part A: Molecular & Integrative Physiology 184:179–186.

Team RC. 2017. R: A language and environment for statistical computing. Vienna: R Foundation for Statistical Computing.

Tonk L, Bongaerts P, Sampayo EM, and Hoegh-Guldberg O. 2013. SymbioGBR: a web-based database of Symbiodinium associated with cnidarian hosts on the Great Barrier Reef. BMC Ecology 13.

Veal CJ, Carmi M, Fine M, and Hoegh-Guldberg O. 2010. Increasing the accuracy of surface area estimation using single wax dipping of coral fragments. Coral Reefs 29:893–897.

Wilson K, Li Y, Whan V, Lehnert S, Byrne K, Moore S, Pongsomboon S, Tassanakajon A, Rosenberg G, Ballment E, Fayazi Z, Swan J, Kenway M, and Benzie J. 2002. Genetic mapping of the black tiger shrimp Penaeus monodon with amplified fragment length polymorphism. Aquaculture 204:297–309.

Wooldridge SA. 2009. Water quality and coral bleaching thresholds: formalising the linkage for the inshore reefs of the Great Barrier Reef, Australia. Marine Pollution Bulletin 58:745–751.

Yellowlees D, Rees T, and Leggat W. 2008. Metabolic interactions between algal symbionts and invertebrate hosts. Plant, Cell & Environment 31:679–694.

## REFERENCES

Franklin, E. C., M. Stat, X. Pochon, H. M. Putnam, and R. D. Gates. 2012. GeoSymbio: a hybrid, cloud based web application of global geospatial bioinformatics and ecoinformatics for Symbiodinium-host symbioses. Molecular Ecology Resources 12:369–373.

Veal, C. J., M. Carmi, M. Fine, and O. Hoegh-Guldberg. 2010. Increasing the accuracy of surface area estimation using single wax dipping of coral fragments. Coral Reefs 29:893–897.

